# Systematic evaluation of transcript normalization methods and quantification pipelines across long-read sequencing platforms

**DOI:** 10.64898/2026.10.10.757893

**Authors:** Alejandro Paniagua, Fabian Jetzinger, Sonia Tarazona, Ana Conesa

## Abstract

Long-read RNA sequencing (lrRNA-seq) has transformed transcriptomics by enabling full-length isoform resolution, yet transcript quantification remains confounded by platform-dependent length biases and tool choices. While a quantification normalization step is common in short-read RNA-seq, this has not yet been extended to lrRNA-seq and the field lacks a systematic analysis of normalization strategies applied to long reads data. Here, we systematically benchmarked quantification pipelines and downstream normalization strategies across Pacific Biosciences (PacBio) and Oxford Nanopore Technologies (ONT) platforms. Using synthetic spike-in controls (SIRVs and ERCC) alongside matched Illumina short-read benchmarks across mouse tissues and human cell models, we evaluated five normalization methods (CPM, TPM, CQN, EDASeq, and Scaled Median Ratio) across five quantification workflows (Bambu, FLAIR, IsoQuant, Oarfish, and Kallisto). Our framework assessed PacBio Iso-Seq and Kinnex protocols together with ONT cDNA and direct RNA (dRNA) sequencing datasets. We demonstrate that standard Transcripts Per Million (TPM) scaling counteracts short-molecule depletion in PacBio datasets, but severely distorts Nanopore expression estimates. For Nanopore libraries, covariate-aware normalizations (CQN and EDASeq) mitigate length bias without TPM’s over-correction penalty. Conversely, relative median ratio normalization (SMR) degrades accuracy and introduces noise in low-variability biological samples. Across computational tools, FLAIR minimized absolute error on short SIRVs, while reference-restricted quantifiers (Oarfish and Kallisto) consistently outperformed transcript reconstruction tools in short-read validation. These findings show that quantification normalization should be applied to lrRNA-seq data and establish platform-specific guidelines for long-read transcriptomics.

## Introduction

Alternative splicing, along with alternative transcription start (TSS) and polyadenylation (poly(A)) site usage, enables single genes to produce multiple structurally distinct RNA isoforms.^1^ Long-read RNA sequencing (lrRNA-seq) has emerged as a powerful approach to resolve this complexity by capturing full-length transcripts in single reads, thereby enabling the direct linkage of distant sequence variables that are lost in fragmented short-read sequencing.^1,2^ This transcript-resolved capability is particularly critical in scenarios involving the identification of cell-type-specific isoform switches during cellular differentiation^1^ or the characterization of aberrant splicing events in human cancers.^3^ Recent theoretical frameworks have mathematically formalized these advantages, proving that longer reads reduce the complexity of transcript deconvolution^4^ and the alignment uncertainty in probabilistic quantification models.^5^ However, in practice, experimental benchmarks highlight platform-specific, length-dependent detection limitations; for instance, Pacific Biosciences (PacBio) bulk sequencing has been shown to under-detect short transcripts (<1.25 kb), whereas Oxford Nanopore Technologies (ONT) bulk runs under-detect long transcripts (>5 kb).^6^ To overcome these limitations, recent computational methods have adopted hybrid approaches that combine the complementary strengths of long and short reads^5,7^ or explicitly model transcript degradation and read quality to improve quantification.^8^ Yet, even with these advanced modeling frameworks, systematic platform-specific biases, including non-linear coverage decay and GC-content distortions, persist in raw datasets. Consequently, choosing appropriate downstream normalization methods remains a critical, yet under-evaluated, step in ensuring accurate abundance estimation across diverse long-read workflows.

The library preparation and biophysical sequencing mechanisms of PacBio and ONT platforms introduce distinct, platform-specific technical biases that complicate abundance estimation.^2^ PacBio Kinnex utilizes circular consensus sequencing (CCS) of concatenated cDNA molecules to yield highly accurate reads, though it remains affected by library-prep size selection, SMRT cell loading bias favoring shorter molecules, and internal priming truncations.^6,9,10^ In contrast, ONT platforms sequence single cDNA or RNA molecules as they translocate through membrane pores, a process sensitive to progressive template degradation and translocation stalling that results in severe, non-linear transcript-body coverage decay.^2,10^ While length and GC biases are well-characterized as linear confounds in short-read sequencing,^11,12^ their non-linear signatures in long-read datasets vary dramatically between platforms,^6,13^ presenting a major challenge for standard normalization approaches.

Standard scaling methods, such as Counts Per Million (CPM) and Transcripts Per Million (TPM), assume linear sequencing probability and fail to correct for the non-linear length biases. More sophisticated joint modeling methods, such as Conditional Quantile Normalization (CQN) and Exploratory Data Analysis (EDASeq), were originally developed to model and remove sample-specific GC-content and length biases in short-read data using generalized additive models or local regressions.^11,12^ EDASeq fits a loess regression of log-counts on transcript length to isolate and remove GC-dependent coverage decay without relying on rigid linear scaling assumptions, offering a flexible normalization strategy alongside CQN’s joint length and GC-content spline corrections.

Here, we present a comprehensive, multi-tool benchmarking framework to evaluate the impact of downstream normalization methods on long-read transcriptome quantification. We developed an automated, Snakemake-based pipeline^14^ to compare five normalization strategies (CPM, TPM, CQN, EDASeq, and ratio-based scaling) across five transcript reconstruction and quantification workflows: Bambu^15^, FLAIR^16^, IsoQuant^17^, Oarfish^18^, and Kallisto^19^. These tools were selected since they represent state-of-the-art, well-established, and actively supported methods that cover diverse algorithmic approaches for isoform quantification. We evaluated them across three diverse studies containing both PacBio and ONT data (**Fig. 1**). Using synthetic spike-ins at known concentrations and orthogonal short-read validation, we demonstrate that no single normalization method is universally applicable across both sequencing platforms. We show that while standard TPM scaling is sufficient and stable for PacBio transcriptomes due to their linear bias profiles, it fails to correct for the non-linear distortions in ONT datasets. Finally, we show that naive ratio-based scaling methods degrade quantification quality across both technologies, establishing clear, platform-specific guidelines for long-read transcriptome abundance normalization.

**Fig. 1.**
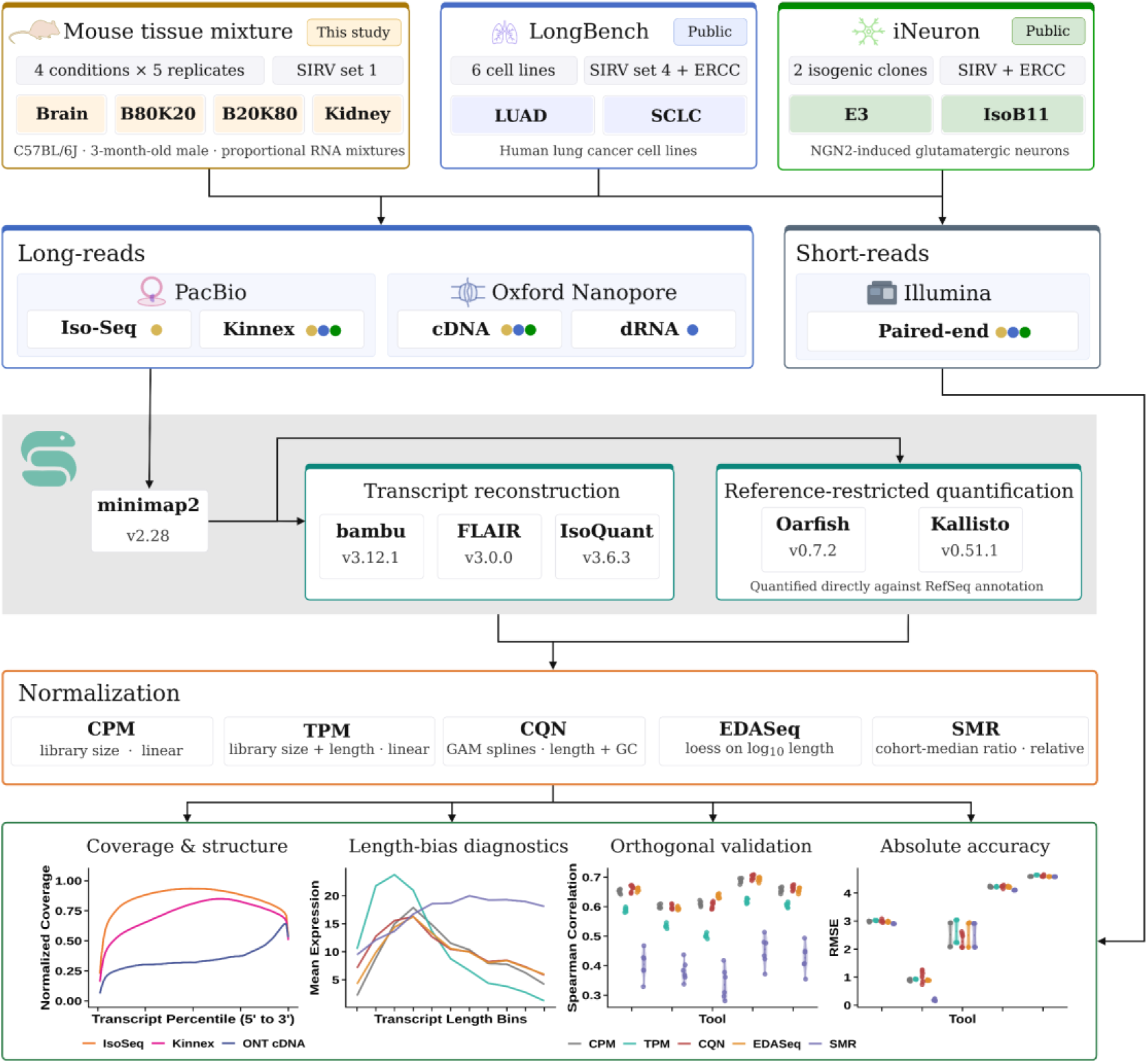
Overview of data collection, processing pipelines, and benchmarking framework. Three benchmark datasets (Mouse tissue mixture, LongBench, and iNeuron) featuring synthetic spike-ins (SIRVs/ERCC) were sequenced using PacBio (Iso-Seq, Kinnex), Oxford Nanopore (cDNA, dRNA), and Illumina technologies. Long-read data was analyzed through transcript reconstruction and reference-restricted quantification tools. Downstream transcript counts were evaluated across five normalization strategies (CPM, TPM, CQN, EDASeq, SMR) based on positional coverage, length-dependent bias, spike-in quantification accuracy (RMSE), and correlation with matched short-read sequencing data.

## Results

### Baseline Sequencing Throughput and Transcript Model Analysis

To establish a baseline for long-read transcriptomic profiling and normalization analyses, we characterized sequencing throughput, read length distributions, and transcript reconstruction properties across three diverse cohorts (**Fig. 2**). The benchmarking cohort comprised a public human lung cancer cell line dataset (LongBench)^13^, a public human induced neurons dataset (iNeuron)^6^, and our mouse tissue mixture dataset derived from brain and kidney.

**Fig. 2.**
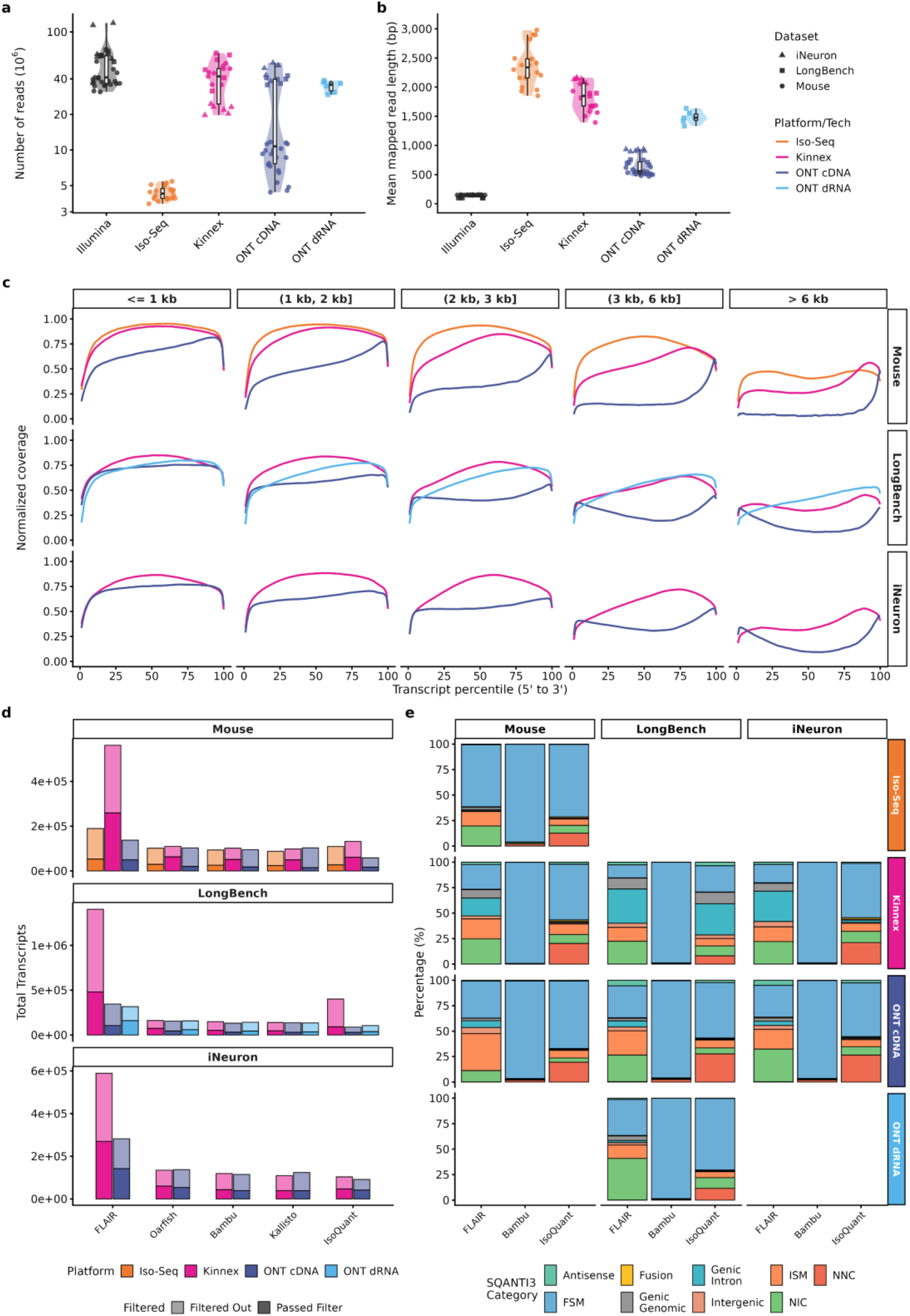
Sequencing throughput, read length distributions, metagene coverage profiles, transcript discovery counts, and SQANTI3 structural classifications across technologies and pipelines. **a**, Library sequencing throughput (number of mapped reads per library, in millions) across sequencing technologies. **b**, Average mapped read length (base pairs) across sequencing technologies. In **a,b**, n = 32 (Illumina), 20 (Iso-Seq), 21 (Kinnex), 32 (ONT cDNA), and 6 (ONT dRNA) samples. Violin plots show the kernel density distribution of values; embedded box plots show the median, interquartile range and whiskers extending to the most extreme values within 1.5× the interquartile range. c, Coverage profiles along the normalized transcript length (5’ to 3’ percentile) across platforms, stratified by transcript length. **d,** Total number of detected transcript models by tool before and after abundance-based filtering across datasets. **e,** Proportion of transcript models classified by SQANTI3 structural categories (Full Splice Match [FSM], Incomplete Splice Match [ISM], Novel in Catalog [NIC], Novel Not in Catalog [NNC], and others) across technologies, discovery tools, and datasets after expression-based filtering.

Sequencing throughput and read lengths varied significantly across platforms, reflecting the technological evolution from PacBio Iso-Seq to Kinnex concatenation and from ONT R9.4.1 to R10.4.1 chemistry (**Fig. 2a,b**). In the mouse tissue cohort, PacBio Kinnex sequencing generated a median of 45.3 million reads per library (range 35.7-54.1 million), representing a ten-fold increase in read count over standard PacBio Iso-Seq runs (median of 4.3 million reads). Conversely, ONT cDNA sequencing using R9.4.1 chemistry yielded a median of 8.6 million reads per library (range 4.4-11.7 million). The R10.4.1 chemistry in the human iNeuron cohort resulted in a substantial throughput expansion, yielding a median of 50.6 million reads for ONT cDNA, compared to 22.5 million reads for Kinnex in the same cohort.

Mapped read length profiles also reflected these platform designs and chemistry evolutions. Standard PacBio Iso-Seq runs recovered the longest median mapped read length of 2,338 bp, whereas the Kinnex concatenation protocol resulted in a median read length of 1,677 bp. ONT cDNA sequencing yielded shorter molecules, with a median mapped read length of 530 bp on the R9.4.1 chemistry. LongBench cell lines runs showed similar throughput profiles for ONT cDNA and Kinnex, while ONT dRNA runs (Kit 14 chemistry) yielded a median of 35.7 million reads pooled from two sequencing runs and a median mapped read length of 1,477 bp. Differences in read length among Iso-Seq, Kinnex, and dRNA are notable compared to ONT cDNA, alongside the expected contrast with Illumina data.

Metagene coverage analysis revealed distinct length-dependent technical decay across platforms (**Fig. 2c**). For reference transcripts under 1 kb, all sequencing technologies exhibited stable, uniform coverage profiles. The sole exception was ONT cDNA on the mouse tissue dataset (R9 chemistry), which showed a rapid decline in coverage towards the 5’ end, resulting in a pronounced 3’ bias. Across all length groups, ONT cDNA showed the most rapid decline in coverage. Conversely, PacBio Kinnex and Iso-Seq profiles exhibited stable average coverage from the 5’ to the 3’ end for short transcripts. However, for longer transcripts, average coverage profiles diverged by library preparation. PacBio and ONT cDNA libraries showed a decrease in average coverage with a slightly bimodal distribution and a more pronounced 3’ hump. In contrast, ONT dRNA libraries exhibited monotonic decay from the 3’ to the 5’ end.

Hierarchical clustering of individual transcript coverage profiles resolved these patterns (**Supplementary Figs. 1–3**). This analysis revealed that the decreased average coverage represented a mixture of partially covered molecules rather than a uniform, intermediate signal across all transcripts. For PacBio and ONT cDNA, distinct clusters of long transcripts showed coverage restricted to either the 5’ or the 3’ region. This indicates that long-read alignments frequently capture partial fragments. In contrast, individual ONT dRNA reads did not show this bimodal division, but instead exhibited a strict 3’ bias. This is biophysically consistent with direct RNA sequencing, which initiates at the 3’ poly(A) tail and proceeds towards the 5’ end. Consequently, premature termination or RNA fragmentation selectively preserves the 3’ region. These coverage biases and transcript truncations were highly reproducible, systematic properties of the platforms rather than stochastic detection artifacts.

We then evaluated how different computational pipelines translate genomic alignments into quantified transcript models. This allowed us to bridge physical read characteristics with downstream abundance quantification, bias diagnostics, and statistical normalization. We executed two main computational strategies: discovery-based workflows (Bambu^15^, FLAIR^16^, and IsoQuant^17^), for which we reconstructed transcript models within individual conditions and merged the resulting transcriptomes to generate a cohort-wide consensus, and reference-restricted quantification workflows (Oarfish^18^ and Kallisto^19^), which estimate expression levels directly against the reference annotation. We assessed the total number of transcripts before and after expression-based filtering, retaining only models that met the minimum abundance and replicate consistency thresholds. (**Fig. 2d**)

Across the evaluated datasets, Kinnex consistently yielded the highest number of discovered transcripts, with the sole exception of the iNeuron cohort. In this dataset, ONT cDNA achieved comparable results, driven by an ONT cDNA read count that was double that of Kinnex (47.9 million versus 22.4 million reads per sample). Among discovery tools, FLAIR exhibited the least stringent filtering, generating high initial transcript counts, such as 562,016 unfiltered (259,445 filtered) models in Mouse Kinnex and 137,460 unfiltered (49,735 filtered) models in Mouse ONT cDNA. By contrast, Bambu (51,268 retained Kinnex; 18,081 retained ONT cDNA) and IsoQuant (61,444 retained Kinnex; 17,071 retained ONT cDNA) remained far more conservative, producing post-filtering model counts that closely aligned with reference-restricted quantifiers like Oarfish (63,184 retained) and Kallisto (49,700 retained; **Fig. 2d**).

SQANTI3^20^ structural classification revealed divergent novel isoform discovery profiles across tools (**Fig. 2e** and **Supplementary Fig. 4**). Prior to filtering, FLAIR and IsoQuant were highly permissive, generating large proportions of novel models (85.97% and 59.64% in Mouse Kinnex, respectively), whereas Bambu was extremely conservative (99.48% Full Splice Matches [FSMs]). Expression-based filtering shifted these proportions in opposite directions: the proportion of novel transcripts discovered by Bambu increased (e.g., 0.52% to 0.96% in Mouse Kinnex), showing its model-based filters selected robust, expressed candidates. Conversely, novel fractions for FLAIR (85.97% to 75.81%) and IsoQuant (59.64% to 45.38%) decreased substantially, indicating that their permissive pipelines produce low-expression, likely spurious isoforms that are systematically removed by expression cutoffs, a dynamic replicated across all cohorts (**Supplementary Fig. 5**).

Splicing model consistency, assessed via unique intron chain (UIC) overlaps, demonstrated a universal core consensus shared across technologies (**Supplementary Fig. 6**). However, tool-and platform-exclusive subsets often exceeded this core. FLAIR combined with Kinnex generated a particularly large set of exclusive transcripts undetected by other tools or platforms. While Kinnex-exclusive subsets were prominent in mouse tissue, they decreased in high-depth iNeuron and LongBench datasets, reflecting improved cross-technology agreement at higher sequencing coverage. Notably, in LongBench, the Kinnex + dRNA exclusive intersection surpassed the combined ONT (dRNA + cDNA) set, highlighting strong concordance between PacBio Kinnex and ONT direct RNA sequencing.

In summary, the collected datasets and analytical methods encompass a wide range of biases, read lengths, sequencing throughput, and transcript reconstruction strategies, providing a comprehensive representation of the lrRNA-seq landscape and recapitulating known properties of the technologies.^6,13,21–23^ Together, they constitute a suitable testbed for evaluating transcript quantification normalization methods.

### Effect of Normalization Methods on Platform-Specific Transcript Length Biases

The systematic differences in read lengths, metagene coverage profiles, and transcript truncations detailed in the previous section directly influence how computational tools reconstruct and quantify transcripts. Because long-read libraries contain a mixture of full-length molecules and truncated fragments, ranging from PacBio Iso-Seq reads (median 2,338 bp) to shorter ONT cDNA fragments (median 530 bp), upstream processing tools must reconcile these partial alignments during expression estimation. Profiling average expression across transcript-length bins revealed that these computational assignments interact with platform-specific library properties to generate divergent transcript length bias signatures across all three benchmarking datasets (**Fig. 3**). Specifically, PacBio libraries (Kinnex and Iso-Seq) exhibited non-linear, unimodal bias profiles, with mean expression peaking at intermediate lengths (around 1.5–2 kb) and declining for both short and very long transcripts. In contrast, ONT cDNA and dRNA libraries displayed severe length-dependent decay.

**Fig. 3.**
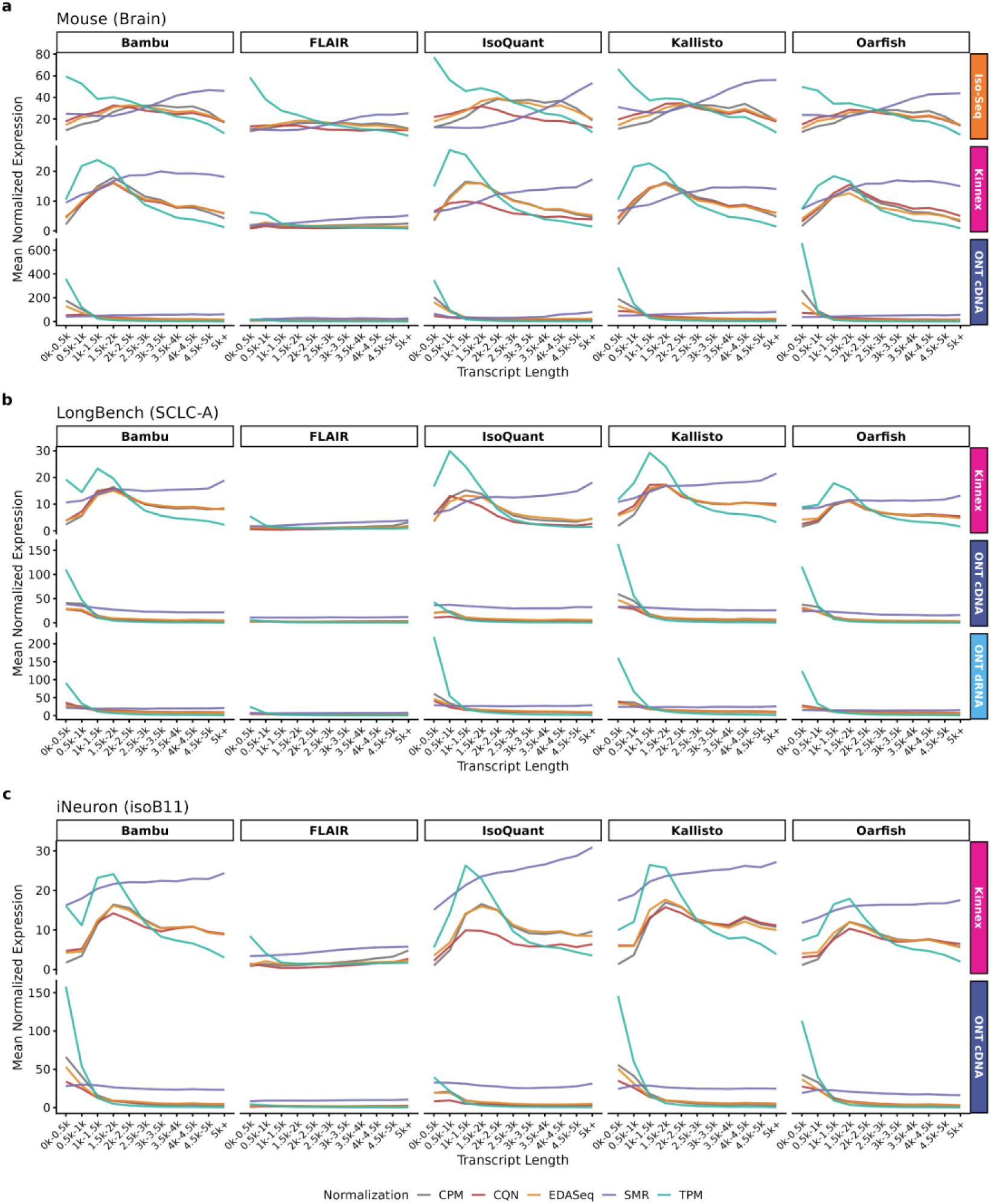
Transcript length-expression relationship under different normalizations. Mean normalized expression across transcript length bins for five normalization methods: CPM, TPM, CQN, EDASeq, and Scaled Median Ratio (SMR) using different tools for quantification. Panels represent different datasets: **a,** mouse Brain; **b,** LongBench (SCLC-A); and **c,** human iNeuron (isoB11).

The choice of downstream normalization method substantially alters these length-dependent profiles (**Fig. 3**). Standard linear-scaling approaches (CPM and TPM) failed to resolve the length-dependent decay in ONT libraries. Because TPM divides raw counts by transcript length, applying it to ONT libraries further increased the abundance of short transcripts that this technology already captures at high rates. In contrast, standard TPM scaling counteracted the depletion of short molecules in PacBio libraries, thereby recovering higher average expression levels for short transcripts. We observed that covariate-aware normalization methods (EDASeq and CQN) did not fully flatten length-expression profiles. Although designed to regress out length-dependent technical effects, both methods left a residual downward trend where average expression levels decayed across transcript length in ONT libraries. Nevertheless, these methods achieved a certain level of mitigation across the transcript length spectrum, as measured by the coefficient of variation of average expression across length bins. In general, CQN and EDASeq successfully reduced the length-expression relationship compared to unadjusted CPM and TPM scaling (**Supplementary Fig. 7**).

In contrast to these technical correction attempts, Scaled Median Ratio (SMR) normalization did not aim to flatten the length-dependent profiles, but rather preserved and resolved the endogenous biological differences between tissues. This biological resolution was most clearly highlighted in the mouse dataset when comparing brain and kidney tissues (**Supplementary Fig. 8**). While technical normalizations (CQN and EDASeq) try to remove length-dependent trends to restore a flat baseline, SMR successfully isolated tissue-specific behaviors. Specifically, brain tissue exhibited a steady, positive length trend where longer transcripts were systematically enriched (**Fig. 3** and **Supplementary Fig. 8**), consistent with the well-established characteristic of mammalian brain tissues expressing exceptionally long transcripts and isoforms.^24^ Conversely, kidney tissue exhibited a systematic negative length trend, reflecting a lower relative abundance of long transcripts (**Supplementary Figs. 8** and **9**). Importantly, this symmetrical inversion, characterized by climbing trends in brain versus decaying trends in kidney, was highly concordant across all sequencing platforms (PacBio Iso-Seq and Kinnex, ONT cDNA) and computational pipelines.

From these analyses, we concluded that different normalization methods substantially alter the relationship between expression quantification and transcript length in lrRNA-seq data. This effect is partly mediated by the transcript reconstruction method used, while the observed patterns are broadly consistent across data types, defined by the combination of sequencing platform and library preparation method. These findings suggest that the most appropriate normalization strategy may not be universal, but rather depends on upstream methodological choices.

### Evaluation of Absolute Quantification Accuracy Using Synthetic Spike-ins

To evaluate absolute abundance recovery and detection sensitivity across long-read pipelines, including normalization methods, we analyzed synthetic spike-in control datasets. Specifically, Spike-in RNA Variants (SIRVs), which mimic complex splicing isoforms, were present in all three cohorts, while External RNA Controls Consortium (ERCC) controls, representing a broad dynamic range of single-isoform transcripts, were added to the public LongBench and iNeuron cohorts. We evaluated absolute quantification accuracy by calculating the root-mean-square error (RMSE) between expected molar concentrations and observed normalized values. In addition, we characterized detection sensitivity as the proportion of spike-in models successfully recovered by each pipeline, resolving distinct performance characteristics across platforms and workflows.

Our analysis of the SIRVs confirmed that absolute quantification accuracy depended strongly on both the sequencing technology and the choice of normalization method (**Fig. 4**). Across all platforms and cohorts, Scaled Median Ratio (SMR) normalization consistently achieved the lowest quantification error. Because SIRVs represent well-defined multi-isoform modules designed with strict stoichiometric relative abundances across sample mixes, ratio-based scaling against a cohort-wide median reference effectively stabilized isoform-level proportions and canceled out shared technical capture artifacts. Among standalone, non-relative normalization methods, performance diverged substantially among sequencing technologies and normalization methods. In the Mouse cohort, TPM normalization minimized PacBio quantification error among standard methods, yielding a mean RMSE of 4.5962 on Iso-Seq and 3.4451 on Kinnex, outperforming CPM (4.8792 on Iso-Seq; 3.8809 on Kinnex), EDASeq (4.8264 on Iso-Seq; 3.7661 on Kinnex), and CQN (4.7032 on Iso-Seq; 3.7591 on Kinnex). Conversely, TPM performed poorly when applied to ONT cDNA data, yielding a mean RMSE of 2.8855, whereas covariate-aware methods (EDASeq: 2.5561; CQN: 2.4761) minimized quantification error. These trends were highly consistent in the public human cohorts. In the iNeuron dataset, TPM minimized PacBio Kinnex error among standard methods (mean RMSE of 4.8198) but collapsed on ONT cDNA (3.1075), where covariate-aware normalizations (EDASeq: 2.7482; CQN: 2.7002) performed best. Similarly, in the LongBench cohort, PacBio Kinnex standard normalizations yielded mean RMSE values of 5.3569 (CPM), 5.267 (TPM), 5.3295 (EDASeq), and 5.3223 (CQN). In ONT cDNA and dRNA runs from the LongBench dataset, standard TPM scaling increased the error to 4.0008 (cDNA) and 3.3814 (dRNA), whereas covariate-aware normalizations successfully reduced the errors (cDNA: 3.3632 for EDASeq, 3.3796 for CQN; dRNA: 3.0334 for EDASeq, 3.0069 for CQN; **Fig. 4**).

**Fig. 4.**
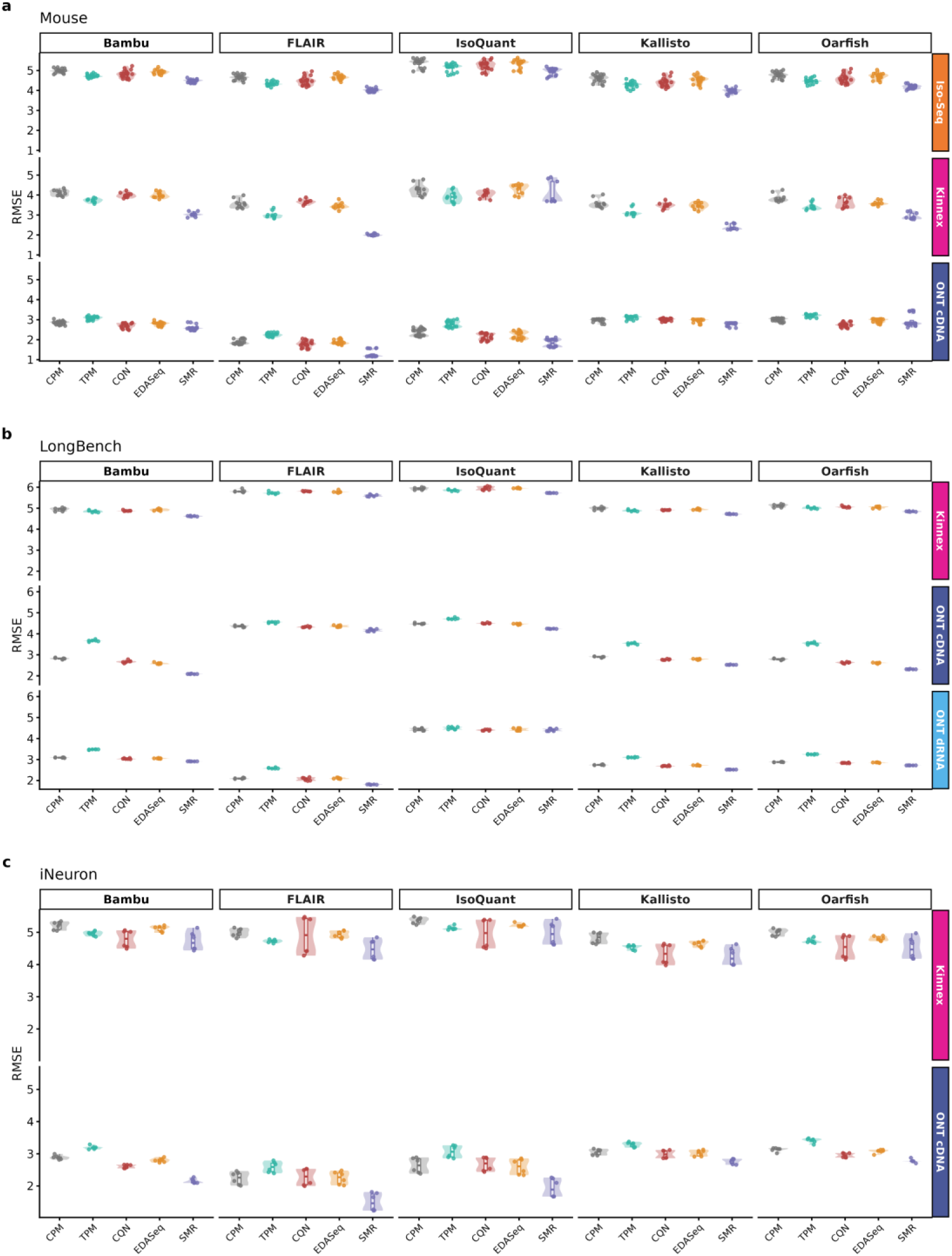
Absolute quantification error of SIRVs. Root-Mean-Square Error (RMSE) between expected molar concentrations and observed normalized values of Spike-in RNA Variants (SIRVs) across upstream tools, normalization methods and sequencing platforms. Panels show different datasets: **a**, mouse (*n* = 49 samples: 20 Iso-Seq, 9 Kinnex and 20 ONT cDNA); **b**, LongBench (*n* = 18: 6 Kinnex, 6 ONT cDNA and 6 ONT dRNA); **c**, human iNeuron (*n* = 12: 6 Kinnex and 6 ONT cDNA). Violin plots show the kernel density distribution of values; embedded box plots show the median, interquartile range and whiskers extending to the most extreme values within 1.5× the interquartile range.

Across tools (excluding SMR), reference-restricted quantifiers consistently recovered abundances with low error across all cohorts, while the transcript reconstruction tools exhibited more divergent behaviors. In our Mouse dataset, FLAIR achieved the lowest error (mean RMSE 3.2735), followed by Kallisto (3.6769), Oarfish (3.7703), Bambu (3.8745), and IsoQuant (3.8824). In the iNeuron cohort, FLAIR again achieved the lowest error (3.6144), followed by Kallisto (3.8351), Bambu (3.9460), Oarfish (3.9582), and IsoQuant (3.9586). However, this pattern shifted in the public LongBench dataset, where reference-restricted quantifiers led, with Kallisto (3.5816) and Oarfish (3.6345) minimizing error, followed by Bambu (3.6672), while FLAIR (4.1319) and IsoQuant (4.9681) lagged.

This shift in tool performance was driven by the presence of exceptionally long SIRVs transcripts (up to 12 kb) in the LongBench cohort **(Supplementary Fig. 10)**, which interacted with platform- and tool-specific length biases across transcript length quantiles (**Supplementary Fig. 11**). While FLAIR and IsoQuant achieved robust performance on short SIRVs when paired with ONT cDNA and dRNA data, they struggled severely with the very long SIRVs transcripts present in the LongBench dataset. In contrast, reference-restricted quantifiers (Kallisto and Oarfish) and the annotation-reliant Bambu recovered these exceptionally long molecules with low error. Across cohorts, PacBio platforms exhibited high quantification errors for the shortest transcript group. This reflects capture depletion from bead-based size selection, which shifted read lengths toward longer molecules compared to the short SIRVs reference distribution (**Supplementary Fig. 10**). Standard TPM scaling was insufficient to correct this shift in PacBio data and remained highly detrimental when applied to ONT libraries. For PacBio datasets, quantification error decreased with transcript length, reaching a minimum in the third length quantile before increasing slightly in the fourth quantile (representing the longest molecules). Conversely, ONT cDNA libraries showed a read length distribution much closer to that of the short SIRVs (**Supplementary Fig. 10**). For these libraries, error remained relatively constant across the length spectrum, with only a slight increase for the longest molecules.

To evaluate how these accuracy patterns extend to datasets with a wider dynamic range, we analyzed the ERCC spike-in controls in the public LongBench and iNeuron datasets (**Supplementary Fig. 12**). We observed a stark contrast in performance compared to the SIRVs analysis, with higher quantification errors for ERCC. Most notably, SMR normalization collapsed when applied to ERCC spike-ins, yielding extremely high errors (RMSE >7.2 across all platforms). This collapse highlighted the differing designs of the two control sets: while SIRVs consisted of distinct groups of overlapping transcript isoforms with large fold changes between groups specifically designed to test splicing complexity, ERCC represented a continuous, broad dynamic range spanning six orders of magnitude of single-isoform transcripts. Standard normalizations remained stable, with TPM, CQN, and EDASeq yielding mean RMSE values of ∼3.23–3.33 on Kinnex and ∼2.86–3.51 on ONT cDNA. Among processing tools, reference-restricted methods performed best, with Oarfish, IsoQuant, and Kallisto achieving mean RMSE values of 3.2723, 3.2819, and 3.2979 in LongBench, respectively, whereas Bambu (3.8526) and FLAIR (4.3319) exhibited higher errors. In the iNeuron cohort, Oarfish and Kallisto led with mean RMSE values of 2.4764 and 2.4778, while FLAIR lagged at 4.4239.

This difference in absolute quantification accuracy revealed by SIRVs and ERCC was consistent with a severe collapse in detection sensitivity for ERCC single-isoforms (dropping to 0%–47% across cohorts) compared to the robust recovery of SIRV transcripts (63%–98% across Mouse and public datasets), particularly under de novo reconstruction pipelines (**Supplementary Fig. 13**).

Together, these results demonstrate that optimal long-read quantification requires aligning normalization methods with platform-specific biases and experimental dynamic range. While ratio-based scaling (SMR) excels in structured isoform modules with fixed stoichiometric proportions, absolute quantification across broad dynamic ranges relies on platform-tailored normalizations, in particular TPM for PacBio and covariate-aware methods for ONT libraries. For computational workflow selection, reference-restricted quantifiers consistently outperform transcript reconstruction methods in both detection sensitivity and quantification stability across extreme transcript lengths.

### Benchmarking Long-Read Normalization Strategies Against Matched Short Reads

Comparison against matched Illumina short-read RNA-seq is commonly used as an orthogonal benchmark of quantification accuracy. We therefore evaluated long-read transcript- and gene-level abundances against matched Illumina short-read data across all cohort samples (**Fig. 5**). Different normalization methods were applied to the long-read data, whereas short-read data were normalized using TPM.^13,18,25^ Because upstream tool choice, downstream normalization method, and sequencing platform operate concurrently and may exhibit complex interdependencies, we decoupled their individual and joint contributions by fitting a linear mixed model (LMM) to the Fisher-*z*-transformed transcript-level correlation coefficients, with biological replicates modeled as random intercepts. Downstream contrasts between factor levels were derived from the fitted models as differences in estimated marginal means (Δz′) using emmeans.^26^

**Fig. 5.**
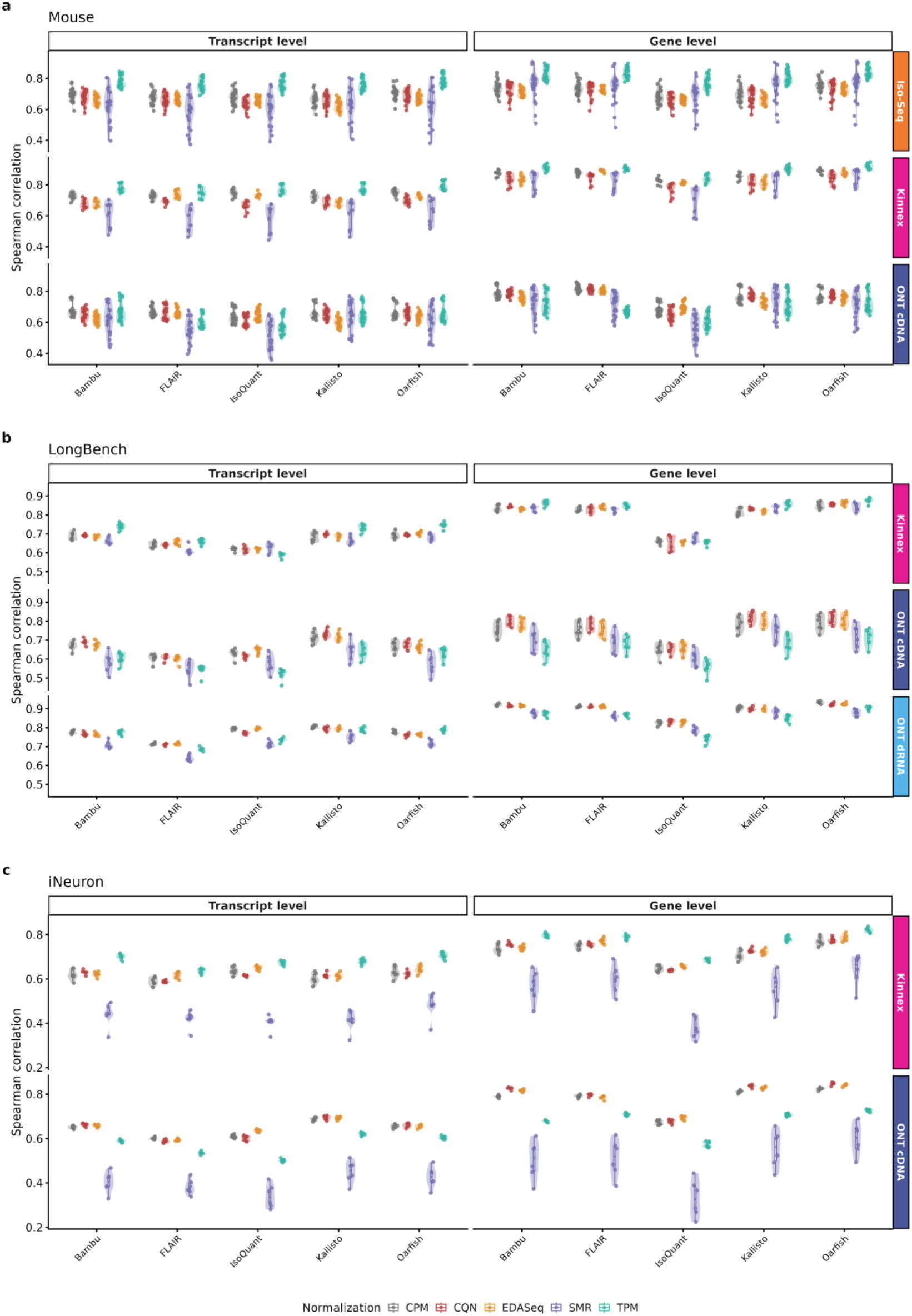
Orthogonal validation correlations against matched Illumina short-read benchmarks. Spearman rank correlation coefficients between long-read expression estimates and matched Illumina short-read data across computational tools and normalization methods. Panels show different datasets: **a**, mouse (*n* = 49 samples: 20 Iso-Seq, 9 Kinnex and 20 ONT cDNA); **b**, LongBench (*n* = 18: 6 Kinnex, 6 ONT cDNA and 6 ONT dRNA); **c**, human iNeuron (*n* = 12: 6 Kinnex and 6 ONT cDNA). Violin plots show the kernel density distribution of values; embedded box plots show the median, interquartile range and whiskers extending to the most extreme values within 1.5× the interquartile range.

Type III Analysis of Variance (ANOVA) on fixed effects revealed that all main effects and their two-way interaction terms were highly significant across all benchmarking cohorts (*p* < 10^−6^; **Supplementary Table 1**). Fixed effects in the LMM explained most of the observed variance in short-read agreement (marginal *R*^2^ = 0.522 in Mouse, 0.944 in iNeuron, and 0.908 in LongBench; conditional *R*^2^ > 0.723, **Supplementary Table 1**). Normalization emerged as a major determinant of correlation and showed a highly significant interaction with sequencing platform in all three cohorts (p < 10⁻⁷⁴; **Supplementary Table 1**), indicating that no single normalization strategy is universally optimal across long-read platforms (**Supplementary Fig. 14**).

Specifically, the model revealed a strong, statistically significant positive synergy between PacBio sequencing chemistry and standard TPM normalization (*Δz*′ = +0.141 for Kinnex in Mouse and *Δz*′ = +0.236 for Iso-Seq in Mouse; *p* < 10^−11^; **Supplementary Figs. 14** and **15**). In PacBio Kinnex runs, gene-level correlations were maximized by TPM, yielding an overall cross-cohort sample median of 0.858 (compared to 0.828 for CPM, 0.820 for EDASeq, and 0.804 for CQN). At the transcript level, TPM reached a median of 0.734, outperforming CPM (0.688), EDASeq (0.674), and CQN (0.661). Similarly, in PacBio Iso-Seq runs, TPM maximized both gene-level (0.816 vs. 0.723 for CPM) and transcript-level correlations (0.766 vs. 0.684 for CPM).

In contrast, the LMM demonstrated that TPM scaling was consistently detrimental when applied to ONT cDNA datasets, significantly reducing agreement relative to CPM in all three cohorts (*Δz*′ = −0.048 in Mouse, −0.112 in iNeuron, and −0.116 in LongBench; p < 10⁻⁴; **Supplementary Table 2**). Covariate-aware normalization methods (CQN and EDASeq) did not manage to substantially outperform CPM (CQN vs. CPM: *Δz*′ = −0.035 to +0.014; EDASeq vs. CPM: *Δz*′ = −0.057 to +0.010, **Supplementary Table 2**). At the gene level, CQN achieved an overall median correlation of 0.784, representing a marginal increase over standard CPM (0.770), while EDASeq (0.763) performed slightly below CPM, and TPM lagged severely at 0.681. At the transcript level, simple CPM led overall with median correlation of 0.654, followed closely by CQN (0.648) and EDASeq (0.642), while TPM lagged at 0.609. Focused interaction plots confirmed CQN and EDASeq performance on ONT cDNA closely parallels simple CPM scaling without providing large additional gains over library-size normalization (**Supplementary Fig. 16**).

Additionally, across all sequencing platforms and cohorts, the model confirmed that relative Scaled Median Ratio (SMR) normalization, where long-read and short-read counts were both transformed into sample-to-reference ratios before correlation computation, yielded lower Spearman correlations than standard absolute scaling methods (SMR vs. CPM: *Δz*′ = −0.029 to −0.336; p < 10⁻³, **Supplementary Table 2**). In PacBio Kinnex runs, the median transcript-level correlation was 0.604 under SMR versus 0.734 under TPM, whereas in ONT cDNA libraries, it was 0.565 under SMR compared to 0.654 under CPM. This decrease was particularly pronounced in the isogenic iNeuron cohort, where biological variance between conditions was limited.

Based on these results, we evaluated tool-level performance under platform-recommended normalizations (TPM for Pacbio and CPM for ONT). Pure reference-restricted quantifiers consistently outperformed transcript reconstruction pipelines across cohorts (*Δz*′ = +0.033 to +0.131 in seven of eight cohort–platform combinations, p < 0.003; Mouse cDNA ONT: *Δz*′ = +0.017, p = 0.13, **Supplementary Table 3**). On PacBio platforms under TPM normalization, Oarfish achieved the highest transcript-level correlations across all four PacBio evaluations (0.789 in Mouse Kinnex, 0.780 in Mouse Iso-Seq, 0.745 in LongBench Kinnex, and 0.705 in iNeuron Kinnex). On ONT cDNA datasets under CPM normalization, Kallisto achieved the highest transcript-level correlations in two of the three cohorts (0.713 in LongBench and 0.689 in iNeuron), both sequenced with the newer R10.4.1 pore chemistry. In contrast, FLAIR and IsoQuant consistently trailed, with significantly negative effects relative to Bambu across all three cohorts (*Δz*′ = −0.021 to −0.113 for FLAIR and −0.037 to −0.070 for IsoQuant; p < 0.01; **Supplementary Table 3**).

Finally, direct platform comparisons and length-stratified evaluations further resolved distinct performance characteristics across sequencing technologies. PacBio Kinnex consistently achieved higher correlation with Illumina than ONT cDNA across both the LongBench and Mouse datasets. In line with TPM’s strong performance on Kinnex data, TPM-normalized Kinnex outperformed ONT cDNA regardless of the normalization method applied to ONT. In the iNeuron dataset, where ONT cDNA throughput doubled that of Kinnex, the gap between platforms narrowed. Notably, ONT dRNA sequencing, which was available only in the LongBench cohort, showed higher correlation than Kinnex (median 0.785 under CPM vs. 0.719 for Kinnex under TPM). Stratifying correlations across transcript length quantiles revealed a systematic decrease in correlation with short reads as transcript length increased. In PacBio datasets, this length-dependent correlation decay occurred primarily for transcripts longer than 4.5 kb and was particularly pronounced for the transcript reconstruction tools. In the iNeuron cohort, we also observed a decrease in correlation for short transcripts. In ONT cDNA libraries, the correlation decay was more immediate, beginning at 1.5 kb. Conversely, dRNA exhibited the most stable correlation behavior across all transcript lengths, showing very limited length-dependent decay in correlation (**Supplementary Fig. 17**).

Altogether, benchmarking against short-read data confirmed that optimal long-read quantification requires a platform-tailored normalization strategy. For PacBio sequencing, linear transcript length division under TPM contributes to counteract the short-transcript depletion introduced by bead-based size selection. In contrast, for ONT cDNA libraries, TPM is actively detrimental, while unadjusted CPM or covariate-aware methods (CQN and EDASeq) are required to preserve quantification fidelity. For tool selection, reference-restricted quantification methods provide improved stability across extreme transcript lengths compared to transcript reconstruction pipelines. Finally, while ratio-based transformations like SMR effectively eliminate technical batch noise across heterogeneous multi-sample cohorts,^27^ they inherently divide out the broad transcriptome-wide dynamic range that drives high absolute rank correlations. In low-variance systems, like the one in the iNeuron cohort, ratio-based scaling isolates low-amplitude relative variations where stochastic measurement noise becomes more prominent.

## Discussion

Our study provides a systematic evaluation of how upstream transcript reconstruction and quantification tools interact with downstream statistical normalizations to shape the accuracy of long-read transcriptomics. By benchmarking these workflows across three diverse datasets, we demonstrate that a single consensus normalization strategy cannot be applied universally. Instead, the distinct biophysical properties of Pacific Biosciences (PacBio) and Oxford Nanopore Technologies (ONT) sequencing dictate platform-specific normalization requirements, resolving long-standing ambiguities in long-read abundance estimation.

This technology-specific performance difference is driven by technical size-selection and translocation biases during library preparation. Standard TPM scaling, which assumes uniform read coverage and linear length decay, effectively counteracts PacBio’s bead-based size selection at the short-molecule end. However, TPM introduces severe distortions when applied to ONT cDNA and dRNA datasets, which exhibit non-linear translocation decay and amplification biases. For these nanopore datasets, covariate-aware regression models (CQN and EDASeq) effectively prevent TPM’s over-correction penalty, however, they fail to substantially outperform unadjusted library-size scaling (CPM).

This limited performance gain occurs because short-read covariate-aware normalizations were developed under the assumption of uniform shotgun fragmentation coverage across reference transcript annotations. In long-read sequencing, we observed two main limitations for the application of this framework: first, long-read platforms display non-uniform positional coverage profiles (such as 5’ truncation gradients and internal priming artifacts); second, individual long reads frequently represent truncated or partial transcript molecules whose physical length distribution deviates significantly from the static reference annotation lengths used as regression covariates. Consequently, regressing against static annotated transcript length introduces model noise, preventing traditional covariate-aware methods from providing substantial gains over unadjusted CPM scaling. Furthermore, the choice of upstream computational tools interacts directly with these normalization curves. Permissive discovery tools like FLAIR generate a high proportion of novel isoforms, which dilutes read allocation and increases downstream quantification uncertainty, whereas reference-restricted quantifiers (Oarfish and Kallisto) and conservative discovery tools (Bambu) preserve stable abundance profiles.

Resolving absolute quantification errors across transcript length bins reveals the limitations of isoform reconstruction tools. While permissive discovery pipelines achieve high detection rates for short transcripts, they struggle with exceptionally long isoforms (up to 12 kb) lacking full-length reads. In contrast, reference-restricted quantifiers and annotation-reliant discovery tools recover long transcripts with low error, as they bypass the splicing-junction assembly phase. These differences are compounded by spatial coverage biases: long transcripts frequently undergo 5’ degradation or internal priming, leading to bimodal coverage profiles that degrade validation correlations with short reads. dRNA sequencing represents a promising alternative, as it bypasses reverse transcription and amplification to yield a more uniform coverage and stable length correlation, though it currently remains constrained by lower throughput compared to cDNA protocols.

Relative normalization methods, such as the Scaled Median Ratio (SMR) framework, exhibit a distinct trade-off between batch harmonization, described by Chen et al.^27^ and absolute accuracy. This transformation systematically reduces direct correlation against validation standards by converting absolute values to relative abundances. This limitation was particularly severe in the highly homogeneous iNeuron cohort, where isogenic cell lines share a stable baseline expression signature. Under SMR, scaling against a cohort median reference mathematically cancels out this biological signal, leaving only low-amplitude stochastic noise that correlates poorly with orthogonal Illumina data. This indicates that ratio-based relative normalizations like SMR can distort absolute abundance profiles when applied to highly homogeneous sample cohorts.

Several technical limitations of our work should be noted. Our validation relies on Illumina short-read datasets as a gold standard, which are themselves subject to multi-mapping ambiguities and GC distortions; and on spike-ins, with limited isoform complexity and isoform length range. Additionally, our dRNA evaluation was restricted to a single dataset, and larger replicated datasets would be needed to confirm these findings. Importantly, our results demonstrate that existing standard normalization methods fail to fully resolve long-read quantification errors, as PacBio and ONT possess distinct technical limitations, such as bead-based size selection versus non-linear translocation decay, that shape fundamentally different biophysical bias spaces. These limitations highlight the need to move beyond empirical scaling frameworks toward explicit mathematical modeling of platform-specific biophysical properties. Precedent for this strategy exists in short-read single-cell RNA-seq, where modeling specific biophysical mechanisms, such as enzyme processivity and kinetics during library preparation, yielded mathematical correction factors that substantially increased transcript quantification accuracy.^28^ Implementing analogous biophysically parameterized correction models for PacBio size-selection profiles or ONT translocation and truncation dynamics represents a promising avenue for refining long-read abundance. These models should also account for the fact that transcript length is coupled with crucial biological processes. For example, multiple studies have shown that age-dependent transcriptional decline and splicing changes occur in a gene-length-biased manner, leading to a global reduction in the transcription of long genes in aged mammalian tissues.^29,30^ Therefore, models should not just merely remove the relation between expression and length.

Based on our findings, we recommend that researchers pair PacBio datasets with standard TPM and reference-restricted quantifiers, adopt CPM or covariate-aware methods (CQN/EDASeq) for Nanopore cDNA and direct RNA to avoid TPM’s severe length-division distortions, and reserve SMR for multi-batch integration where global bias cancellation is required. However, this is a fast-moving field and the technologies are under constant development. For example, Oxford Nanopore made recent improvements in cDNA library preparation claiming longer reads generation, while other community driven efforts also aim in the same direction, with protocols like CapTrap or improvements in retrotranscription. ^31,32^ These are not part of our study as they are not main-stream methodologies yet and we lack replicated data that include spike-ins and several technologies. However, they should be available soon. To update our current recommendations, we provide a Snakemake pipeline to easily analyze newly generated data as technology continues evolving.

## Methods

### Sample acquisition, nucleic acid extraction, and sequencing

#### Mouse tissue proportional mixture dataset

To evaluate transcript-level abundance estimation, we designed a four-condition experimental cohort with five biological replicates per condition, consisting of pure mouse brain, pure mouse kidney, an 80% brain / 20% kidney (B80K20) mixture, and a 20% brain / 80% kidney (B20K80) mixture. Prior to library preparation, Lexogen Spike-in RNA Variants (SIRV) controls were added to the RNA samples: pure brain samples and B80K20 mixtures were spiked with the SIRV E0 mix, pure kidney samples were spiked with the SIRV E1 mix, and B20K80 mixtures were spiked with the SIRV E2 mix. The animal handling, tissue collection, dissection, and RNA extraction procedures for the samples were conducted as described in Jetzinger et al.^33^ Briefly, tissues were collected from 3-month-old male C57BL/6J wild-type mice, followed by intracardial perfusion, specific organ dissection, and total RNA isolation using the Qiagen RNeasy Mini Kit. We expanded the dataset of pure samples by preparing RNA mixes in different proportions.

Libraries were prepared and sequenced across multiple long-read platforms and one orthogonal short-read technology. For Oxford Nanopore Technologies (ONT) cDNA sequencing, barcoded cDNA libraries were synthesized using the Nanopore SQK-PCB111-24 PCR-cDNA Barcoding Kit and sequenced on R9.4.1 flow cells using a PromethION platform across all 20 samples. For PacBio Iso-Seq, cDNA libraries were barcoded using the NEBNext Single Cell/Low Input cDNA Synthesis and Amplification Kit, prepared with the Iso-Seq SMRTbell prep kit 3.0, and sequenced on a PacBio Sequel IIe. Furthermore, to benchmark newer high-throughput long-read sequencing, a subset of 9 samples (3 pure brain, 3 pure kidney, and 3 B20K80 mixture samples) was sequenced using the PacBio Kinnex RNA kit on a PacBio Revio system. For orthogonal validation, matching RNA aliquots from all 20 samples were prepared using the TruSeq Stranded mRNA Library Prep Kit and sequenced on an Illumina NovaSeq platform (2×150bp paired-end reads) at Novogene.

#### Public human lung cancer cell line dataset (LongBench)

The public LongBench^13^ dataset consists of bulk transcriptomic profiling of six human lung cancer cell lines representing two major subtypes: lung adenocarcinoma (LUAD: H1975, HCC827, and H2228) and small cell lung cancer (SCLC: SHP77, H69, and H146). Prior to library preparation, Lexogen Spike-in RNA Variants (SIRVs) set 4 and External RNA Controls Consortium (ERCC) spike-ins were added to the samples. Libraries were prepared and sequenced across bulk PacBio Kinnex sequencing, bulk ONT cDNA sequencing, bulk ONT direct RNA (dRNA) sequencing, and orthogonal Illumina NextSeq2000 paired-end sequencing.

#### Public human iNeuron dataset

The public iNeuron dataset consists of bulk transcriptomic profiling of human NGN2-induced neurons^6^. These glutamatergic neurons were differentiated over a 21-day period from Fragile X Syndrome (FXS) patient-derived induced pluripotent stem cells (iPSCs; patient reporter clone E3) and a CRISPR-edited isogenic rescue clone (IsoB11). Clone E3 is characterized by silencing of the *FMR1* gene, while clone IsoB11 has a rescued *FMR1* locus. Lexogen Spike-in RNA Variants (SIRVs) and External RNA Controls Consortium (ERCC) spike-ins were added to the samples prior to library preparation. Libraries were prepared and sequenced across bulk PacBio Kinnex sequencing, bulk ONT PCR-cDNA sequencing, and orthogonal Illumina NovaSeq paired-end sequencing.

### Metagene coverage analysis

To characterize technical coverage decay and artifact distributions across transcript bodies, we extracted read coverage profiles across 100 percentile bins (from 5’ to 3’) for each transcript. Long reads were aligned to reference transcriptome sequences (NCBI RefSeq GCF_000001405.40 for human, GCF_000001635.27 for mouse) using minimap2 (v.2.28)^34^ with presets *-x map-hifi* (for PacBio) or *-x map-ont* (for ONT). Base-resolution depth was computed using samtools depth (v.1.19)^35^. Coverage values were aggregated into 100 bins per transcript for all models with spliced length longer than 100bp and a minimum coverage of 10 reads in at least one bin. Individual transcript profiles were normalized relative to their maximum bin value to scale coverage from 0 to 1.

For each dataset, the normalized coverage profiles were averaged across samples for each transcript, and transcripts were stratified by length into different size groups. To resolve sub-populations of partial molecules and template fragmentation, transcript profiles within each size group were clustered using agglomerative hierarchical clustering with Euclidean distance and Ward’s minimum variance linkage.

### Obtaining transcripts from lrRNA sequencing data

#### Long-read transcriptome reconstruction and quantification pipeline

We developed an automated, modular Snakemake workflow to process, reconstruct, and quantify long-read sequencing data.^14^ The pipeline handles raw fastq and BAM inputs, manages tool dependencies through isolated Conda environments, and supports parallel execution on SLURM clusters.

Long-read genomic alignments were generated using minimap2 (v.2.28)^34^ with parameters *-ax splice:hq -uf --MD* (for PacBio Kinnex reads) or preset *-x map-ont* (for ONT cDNA and dRNA reads) against reference genomes (GRCm39 or GRCh38). We benchmarked five long-read transcriptomics tools, categorized into discovery-based and reference-restricted workflows.

#### Discovery-based transcript reconstruction workflows

Discovery-based transcript reconstruction was performed using three workflows. Bambu (v.3.12.1)^15^ was run with default settings. Only expressed transcripts were kept in the final GTF. FLAIR (v.3.0.0)^16^ transcript reconstruction was run through the wrapper function *flair transcriptome* using *--stringent --check_splice* parameters. Sample-level quantification was computed using flair quantify with *--sample_id_only --quality 0 --generate_map* parameters. IsoQuant (v.3.6.3)^17^ was run with the appropriate *–data–type*, nanopore or pacbio, based on the technology used.

#### Reference-restricted transcript quantification workflows

Reference-restricted transcript quantification was performed using two workflows. Oarfish (v.0.7.2)^18^, unlike the previously described tools, requires the alignment of reads against the transcriptome sequence. Reads were aligned to the transcriptome using minimap2 with presets *- x map-hifi* (for PacBio) or *-x map-ont* (for ONT). Reads were quantified using Oarfish with parameters *--filter-group no-filters --model-coverage* to account for coverage bias along the transcript body. Kallisto (v.0.51.1)^19^ index was constructed using the recommended k-mer size 63 for long read quantification. Long reads were pseudoaligned using kallisto bus *--long -- threshold=0.8 -x bulk*, and the resulting BUS file was sorted via bustools sort and counted via *bustools count --cm -m* to yield raw abundance matrices, which were quantified at the transcript level via *kallisto quant-tcc --long -P [PacBio/ONT]*.

#### Multi-sample transcriptome consolidation and reference matching

For discovery-based workflows (Bambu, FLAIR, IsoQuant), reconstructed transcript models must be consolidated across samples and conditions to build a unified transcriptome. Transcript models were merged using Transcriptome Annotation by Modular Algorithms (TAMA)^36^ merge with parameters *-m 0 -d merge_dup -a 50 -z 50* to collapse redundant structures while preserving splice junction boundaries.

The consolidated transcriptome was passed to SQANTI3 (v.4.2)^20^ for structural classification against reference annotations. Reconstructed transcripts were compared against reference annotations (NCBI RefSeq GCF_000001405.40 for human, NCBI RefSeq GCF_000001635.27 for mouse) and classified into major structural categories based on splice junction concordance: Full Splice Match (FSM), where the transcript coordinates match a reference transcript model at all splice junctions; Incomplete Splice Match (ISM), where the transcript matches consecutive reference splice junctions but is truncated at the 5’ or 3’ end; Novel In Catalog (NIC), where the transcript recombines known reference donor and acceptor splice sites in a novel junction combination; and Novel Not in Catalog (NNC), where the transcript contains at least one novel splice site (donor or acceptor) not present in the reference catalog. Finally, we filtered out transcripts that failed to reach at least 5 counts across all biological replicates in at least one experimental condition.

To evaluate changes in the proportion of novel versus known transcripts before and after abundance-based filtering, we categorized transcripts into two groups: Known (FSM) and Novel (comprising all non-FSM structural categories). For each computational workflow and sequencing technology combination, we constructed a 2 × 2 contingency table crossing transcript status (Known vs. Novel) by filtering outcome (Passed Filter vs. Failed Filter). Pearson’s chi-squared test of independence was performed on these tables to evaluate whether the probability of passing the abundance-based filter differed significantly between known and novel transcripts. Resulting p-values were adjusted for multiple hypothesis testing using the Bonferroni correction method.

### Downstream abundance normalization methods

Raw count matrices were normalized using five distinct strategies. To ensure comparability across methods, all resulting abundance matrices were scaled to a uniform library depth of 10⁶ as a final step.

#### CPM (Counts Per Million)

CPM scales raw counts for transcript *i*, *C_i_*_,*j*_, linearly by the total library size of sample *j*, adjusting for differences in sequencing depth without accounting for transcript length:

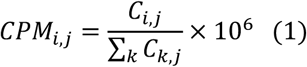

#### TPM (Transcripts Per Million)

TPM scales raw counts linearly by both transcript length *L_i_* and library size, assuming a uniform, linear probability of sequencing coverage across the transcript body:

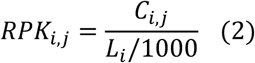

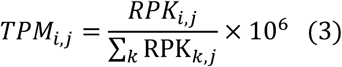

#### Loess-based length bias regression (EDASeq)

EDASeq^12^ normalization is a local regression model designed to regress out non-linear GC bias from raw counts. Instead, we applied it based on transcript length. For each sample, *gcLoess* fits a local regression of log-counts on log₁₀-transformed transcript length, using only transcripts with positive counts falling below the 99th percentile of that sample’s count distribution. The fit used a second-degree local polynomial and a span of 0.75. Each transcript was then scaled by the deviation of the fitted curve from the median log-count of the transcripts used in the fit, yielding length-corrected quantification.

#### Conditional Quantile Normalization (CQN)

CQN^11^ is a quantile regression model designed to regress out non-linear GC and length biases from raw counts. For each sample, *cqn* fits a median regression of log₂ counts on GC content and log₂ transcript length, with log₂ library size as an offset, using only transcripts with mean counts above 50. Both covariates enter as natural cubic splines with 5 knots placed at the 0.025, 0.25, 0.5, 0.75 and 0.975 quantiles, and the coefficients are estimated by median regression. The residuals from this fit, which carry the GC- and length-corrected expression, are then normalized across samples by a modified quantile normalization in which each sample CDF is estimated as a weighted average of its empirical CDF and a Gaussian-mixture fit, so that transcripts outside the fitted subset can also be mapped onto the common target distribution. Values are returned on the log₂ CPM scale and expression estimates were obtained as 2*^yi^*^,*j*+offset^*^i^*^,*j*^.

#### Scaled Median Ratio (SMR)

To assess transcript abundance relative to a cohort-wide baseline while preserving a CPM equivalent scale, we propose Scaled Median Ratio (SMR) normalization. This strategy leverages the stability of relative expression ratios while transforming the outputs back to a non-negative, standard library size scale.

First, raw transcript counts *C_i_*_,*j*_ for transcript *i* and sample *j* are scaled to CPM to account for differences in sequencing depth. For each transcript *i*, the cohort-wide baseline is computed as the median CPM expression across all samples. We then calculate the relative abundance ratio for each sample by dividing the observed CPM by the transcript median, adding a stabilization offset (pseudocount) of 1 to the denominator to prevent division by zero and mitigate high variance in low-abundance transcripts:

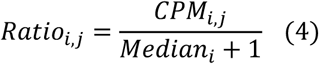

Finally, to convert these ratios into a standard, comparable library size scale and represent them as pseudocounts, we apply CPM normalization to the ratio matrix, scaling the ratios to sum to 10^6^ for each sample:

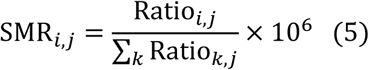

### Transcript length bias diagnosis

To evaluate technical transcript length biases across tool and normalization combinations, we profiled expression levels stratified by transcript length. For all evaluated normalization methods, abundances were analyzed on the CPM scale to ensure uniform library sizes and render the profiles directly comparable. To stratify transcripts into length bins, we computed the trimmed mean of non-zero expression values across biological conditions (excluding the top and bottom 2.5% outliers) for each bin. To measure the flatness of these profiles, we computed the coefficient of variation (CV) across the bins.

### Absolute quantification accuracy and detection sensitivity using synthetic spike-ins

To evaluate absolute abundance recovery, we utilized synthetic spike-in controls. Spike-in RNA Variants (SIRVs) were present in all three cohorts, while External RNA Controls Consortium (ERCC) controls were added to the public LongBench and iNeuron datasets. We evaluated absolute quantification accuracy by calculating the Root Mean Square Error (RMSE) in log scale between known expected molar concentrations and observed normalized values across all combinations of sequencing technologies, normalizations, and processing tools. Additionally, to evaluate transcript detection, we characterized detection sensitivity as the proportion of expected spike-in transcript models successfully recovered by each pipeline.

### Relative abundance validation against short-read benchmarks

Endogenous transcript abundance estimates were validated against matched Illumina short-read profiles. Spearman rank correlation coefficients (*ρ*) were calculated between the long-read estimates and the matching short-read pseudoaligned abundance estimates (quantified using kallisto v0.51.1)^37^ across all conditions. For these comparisons, Illumina counts were normalized using TPM, except when validating long-read estimates normalized by SMR, in which case the corresponding Illumina counts were also normalized using SMR to ensure a matching relative reference frame.

To decouple the concurrent and interacting effects of upstream computational tools, downstream normalization methods, and sequencing technologies on quantification consistency, we fitted Linear Mixed-Effects Models (LMMs) to the transcript-level Spearman correlation coefficients. Correlations were previously transformed into an unbounded, approximately normal scale using Fisher’s *z*-transformation:

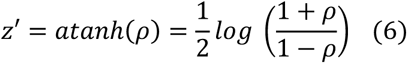

For each cohort independently, the mixed model was formulated using the lme4 (v.1.1-37)^38^ and lmerTest (v.3.2-1)^39^ packages in R (v.4.1.2)^40^. The model included tool, normalization, and technology (sequencing platform) as fixed effects and their two-way interaction terms. Biological replicates were modeled as random intercepts, accounting for the non-independence of measurements taken on the same sample across tools and normalization methods. Baseline reference levels were set to Bambu for tool, CPM for normalization, and ONT cDNA for technology. Degrees of freedom and statistical significance were determined using Type III Analysis of Variance (ANOVA) F-tests using Satterthwaite’s approximation for the denominator degrees of freedom. Marginal coefficients of determination (*R*^2^, variance explained by fixed effects) and conditional coefficients of determination (*R*^2^, total variance explained by fixed and random effects combined) were calculated using the Nakagawa and Schielzeth framework directly from the fitted fixed-effect and random-intercept variance components.^41^

Effects were reported as contrasts between estimated marginal means (EMMs), computed with the emmeans package (v.2.0.2)^26^ using Satterthwaite degrees of freedom and equal weighting across the levels of non-specified factors. Contrasts are expressed as differences on the Fisher z′ scale (*Δz*′). Marginal main effects (each level vs. the reference level, averaged over the remaining factors) and two-way interaction contrasts were adjusted for multiple testing using the Holm method within each term type. Normalization effects within each sequencing platform were evaluated against CPM with Dunnett adjustment and across all pairs of methods with Tukey adjustment. Tool performance was evaluated under the platform-recommended normalization through Tukey-adjusted pairwise comparisons and a single linear contrast comparing the mean of reference-restricted quantifiers (Kallisto and Oarfish) with that of transcript reconstruction pipelines (Bambu, FLAIR and IsoQuant). Statistical significance was set at an adjusted p < 0.05.

## Data availability

The mouse mixture sequencing data generated in this study have been deposited in the European Nucleotide Archive (ENA) under accession PRJEB123244. Raw ONT cDNA, PacBio, and Illumina sequencing data for the pure samples are available under accession numbers PRJEB94912 and PRJEB85167.

Publicly available dataset LongBench analyzed in this study was retrieved from Gene Expression Omnibus (GEO) under accession GSE303762. iNeuron dataset was retrieved from Array Express under accession E-MTAB-16791.

## Code availability

Snakemake pipeline and scripts necessary to reproduce the plots in this paper are available in GitHub (https://github.com/alexpan00/transcript_bias, https://github.com/alexpan00/length_bias_plots). Input data files surpassing GitHub size limitations were uploaded to Zenodo (https://doi.org/10.5281/zenodo.22109031)

## Acknowledgements

A.P. thanks Carolina Monzó for insightful discussions and Tianyuan Liu for help with the design of Fig. 1.

The computations were performed on the high-performance computing cluster Garnatxa at the Institute for Integrative Systems Biology (I^2^SysBio), I^2^SysBio is a mixed research centre formed by University of Valencia (UV) and Spanish National Research Council (CSIC).

## Author Contributions

A.P. developed the main part of the code, performed analyses and drafted the manuscript. F.J. contributed to the development of the code. A.C. and S.T. supervised the study and manuscript drafting. All authors approved the manuscript.

## References

1. Belchikov, N. et al. Understanding isoform expression by pairing long-read sequencing with single-cell and spatial transcriptomics. Genome Res. 34, 1735–1746 (2024).

2. Monzó, C., Liu, T. & Conesa, A. Transcriptomics in the era of long-read sequencing. Nat. Rev. Genet. 26, 681–701 (2025).

3. Anvar, S. Y. et al. Full-length mRNA sequencing uncovers a widespread coupling between transcription initiation and mRNA processing. Genome Biol. 19, (2018).

4. Zhang, B., Guo, Y., Liu, G. & Zou, M. Quantifying transcript complexity via the condition number of gene-specific random matrix. Brief. Bioinform. 27, bbag134 (2026).

5. Li, H. et al. Improving gene isoform quantification with miniQuant. Nature Biotechnology 2025 44:3 44, 477–489 (2025).

6. Botta, G. et al. Benchmarking long-read RNA-seq across modalities, methods, and sequencing depth in iNeurons. bioRxiv 2026.04.01.715783 (2026) doi:10.64898/2026.04.01.715783.

7. Apostolides, M. et al. Accurate isoform quantification by joint short- and long-read RNA-sequencing. bioRxiv https://doi.org/10.1101/2024.07.11.603067 (2024) doi:10.1101/2024.07.11.603067.

8. Hu, Y. et al. LIQA: long-read isoform quantification and analysis. Genome Biol. 22, 182 (2021).

9. Al’Khafaji, A. M. et al. High-throughput RNA isoform sequencing using programmed cDNA concatenation. Nature Biotechnology 2023 42:4 42, 582–586 (2023).

10. Calvo-Roitberg, E., Daniels, R. F. & Pai, A. A. Challenges in identifying mRNA transcript starts and ends from long-read sequencing data. Genome Res. 34, 1719–1734 (2024).

11. Hansen, K. D., Irizarry, R. A. & Wu, Z. Removing technical variability in RNA-seq data using conditional quantile normalization. Biostatistics 13, 204 (2012).

12. Risso, D., Schwartz, K., Sherlock, G. & Dudoit, S. GC-content normalization for RNA-Seq data. BMC Bioinformatics 12, (2011).

13. You, Y., et al. Benchmarking long-read RNA-sequencing technologies with LongBench: a cross-platform reference dataset profiling cancer cell lines with bulk and single-cell approaches. bioRxiv 2025.09.11.675724 (2026) doi:10.1101/2025.09.11.675724.

14. Mölder, F. et al. Sustainable data analysis with Snakemake. F1000Res. 10, 33 (2025).

15. Chen, Y. et al. Context-aware transcript quantification from long-read RNA-seq data with Bambu. Nature Methods 2023 20:8 20, 1187–1195 (2023).

16. Tang, A. D. et al. Full-length transcript characterization of SF3B1 mutation in chronic lymphocytic leukemia reveals downregulation of retained introns. Nature Communications 2020 11:1 11, 1–12 (2020).

17. Prjibelski, A. D. et al. Accurate isoform discovery with IsoQuant using long reads. Nature Biotechnology 2023 41:7 41, 915–918 (2023).

18. Jousheghani, Z. Z., Singh, N. P. & Patro, R. Oarfish: enhanced probabilistic modeling leads to improved accuracy in long read transcriptome quantification. Bioinformatics 41, i304 (2025).

19. Loving, R. K. et al. Long-read sequencing transcriptome quantification with lr-kallisto. PLoS Comput. Biol. 21, e1013692 (2025).

20. Pardo-Palacios, F. J. et al. SQANTI3: curation of long-read transcriptomes for accurate identification of known and novel isoforms. Nature Methods 2024 21:5 21, 793–797 (2024).

21. Pardo-Palacios, F. J. et al. Systematic assessment of long-read RNA-seq methods for transcript identification and quantification. Nat. Methods 21, 1349–1363 (2024).

22. Chen, Y. et al. A systematic benchmark of Nanopore long-read RNA sequencing for transcript-level analysis in human cell lines. Nature Methods 2025 1–12 (2025) doi:10.1038/s41592-025-02623-4.

23. Wissel, D. et al. A systematic benchmark of high-accuracy PacBio long-read RNA sequencing for transcript-level quantification. Genome Biol. https://doi.org/10.1186/S13059-026-03988-1 (2026) doi:10.1186/S13059-026-03988-1.

24. Miura, P., Shenker, S., Andreu-Agullo, C., Westholm, J. O. & Lai, E. C. Widespread and extensive lengthening of 3’ UTRs in the mammalian brain. Genome Res. 23, 812–825 (2013).

25. Dong, X. et al. Benchmarking long-read RNA-sequencing analysis tools using in silico mixtures. Nat. Methods 20, 1810–1821 (2023).

26. Lenth, R. V & Piaskowski, J. emmeans: Estimated Marginal Means, aka Least-Squares Means. https://CRAN.R-project.org/package=emmeans (2026).

27. Chen, Q. et al. A ratio-based framework using Quartet reference materials for integrating long- and short-read RNA-seq. bioRxiv 2025.09.15.676287 (2025) doi:10.1101/2025.09.15.676287.

28. Archer, N., Walsh, M. D., Shahrezaei, V. & Hebenstreit, D. Modeling Enzyme Processivity Reveals that RNA-Seq Libraries Are Biased in Characteristic and Correctable Ways. Cell Syst. 3, 467–479.e12 (2016).

29. Stoeger, T. et al. Aging is associated with a systemic length-associated transcriptome imbalance. Nature Aging 2022 2:12 2, 1191–1206 (2022).

30. Parast, S. et al. Diminished transcriptional activity and splicing changes drive gene length–biased rewiring in the aging transcriptome. Proceedings of the National Academy of Sciences 123, e2607264123 (2026).

31. Carbonell-Sala, S. et al. CapTrap-seq: a platform-agnostic and quantitative approach for high-fidelity full-length RNA sequencing. Nature Communications 2024 15:1 15, 5278-(2024).

32. Chou, C.-L. et al. Single-cell RNA-seq using UltraMarathonRT expands the known transcriptome. bioRxiv 2025.10.06.680646 (2025) doi:10.1101/2025.10.06.680646.

33. Jetzinger, F. et al. Handling biological replicates in long-read RNA sequencing data by joining or not joining. Nature Communications 2026 https://doi.org/10.1038/S41467-026-77367-6 (2026) doi:10.1038/S41467-026-77367-6.

34. Li, H. Minimap2: pairwise alignment for nucleotide sequences. Bioinformatics 34, 3094–3100 (2018).

35. Danecek, P. et al. Twelve years of SAMtools and BCFtools. Gigascience 10, 1–4 (2021).

36. Kuo, R. I. et al. Illuminating the dark side of the human transcriptome with long read transcript sequencing. BMC Genomics 21, 1–22 (2020).

37. Bray, N. L., Pimentel, H., Melsted, P. & Pachter, L. Near-optimal probabilistic RNA-seq quantification. Nat. Biotechnol. 34, 525–527 (2016).

38. Bates, D., Mächler, M., Bolker, B. M. & Walker, S. C. Fitting Linear Mixed-Effects Models using lme4. J. Stat. Softw. 67, (2014).

39. Kuznetsova, A., Brockhoff, P. B. & Christensen, R. H. B. lmerTest Package: Tests in Linear Mixed Effects Models. J. Stat. Softw. 82, 1–26 (2017).

40. R Core Team. R: A Language and Environment for Statistical Computing. https://www.R-project.org/ (2021).

41. Nakagawa, S. & Schielzeth, H. A general and simple method for obtaining R2 from generalized linear mixed-effects models. Methods Ecol. Evol. 4, 133–142 (2013).

