## Supplementary Material for "Systematic evaluation of transcript normalization methods and quantification pipelines across long-read sequencing platforms"

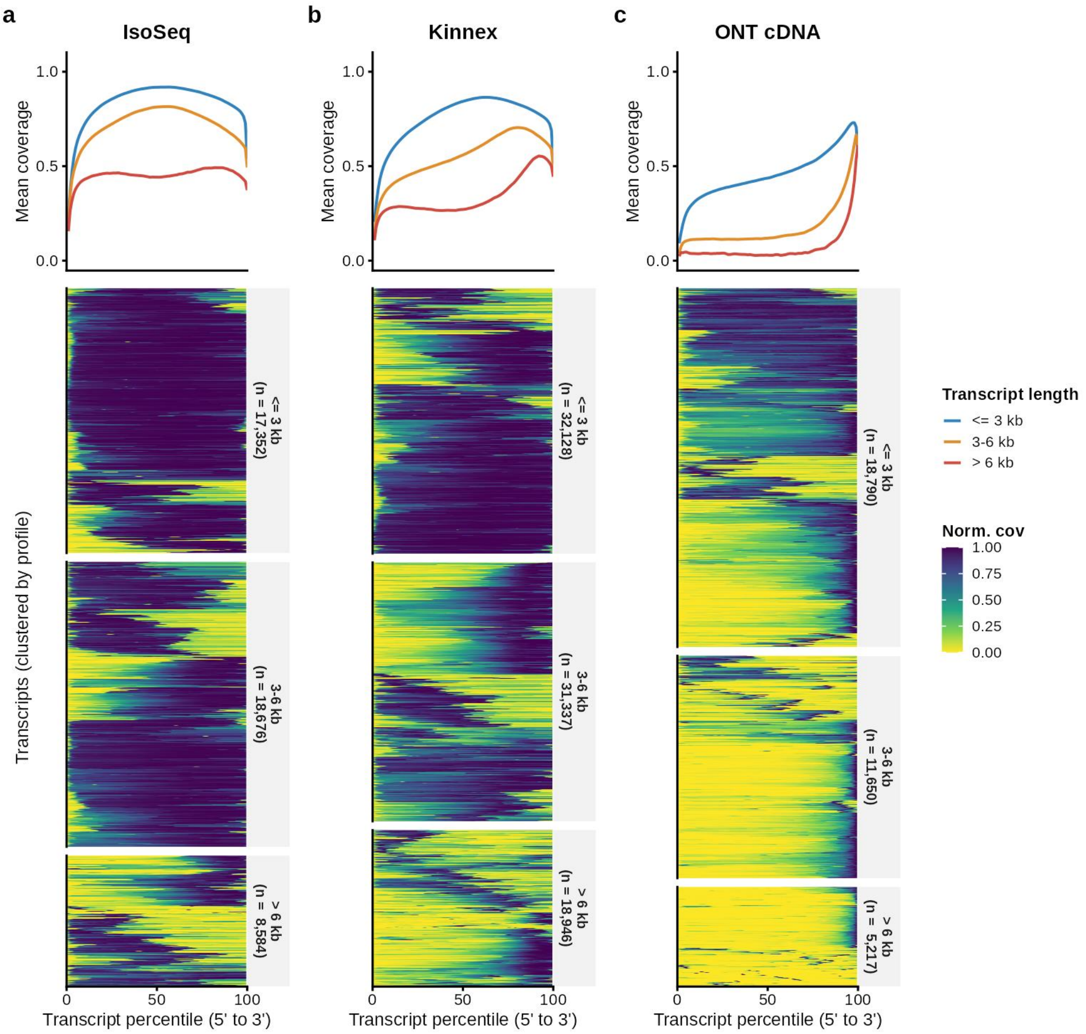


***Supplementary Fig. 1. Hierarchical clustering of individual transcript coverage profiles in the mouse tissue dataset***. Relative read coverage across transcript percentiles for individual transcripts clustered hierarchically. Plots are stratified by sequencing platform (PacBio Iso-Seq, PacBio Kinnex, ONT cDNA) and transcript length bins. Individual rows represent single transcript models.

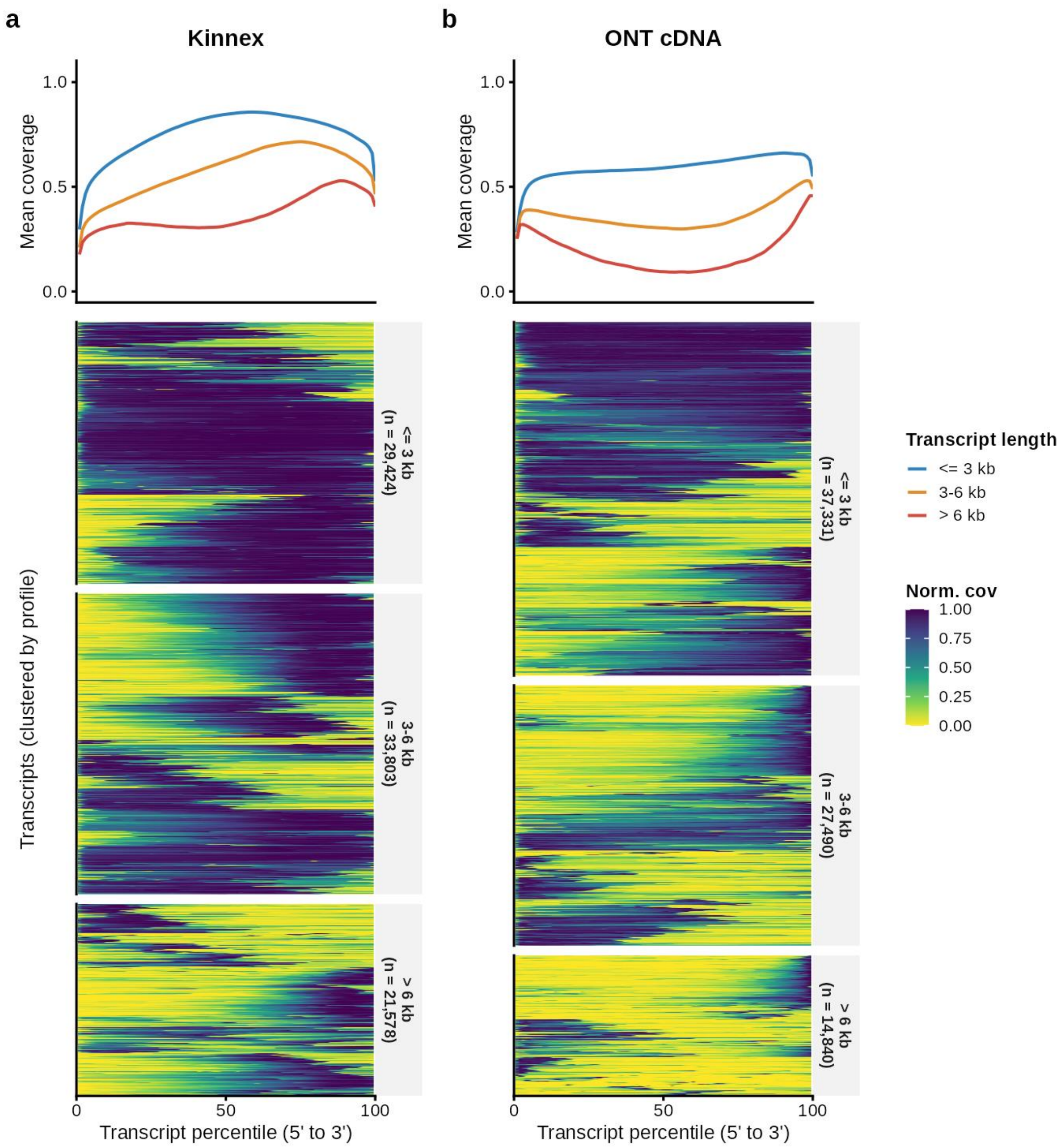


***Supplementary Fig. 2. Hierarchical clustering of individual transcript coverage profiles in the human iNeuron dataset***. Relative read coverage across transcript percentiles for individual transcripts clustered hierarchically across PacBio Kinnex and ONT cDNA technologies, stratified by transcript length bins. Individual rows represent single transcript models.

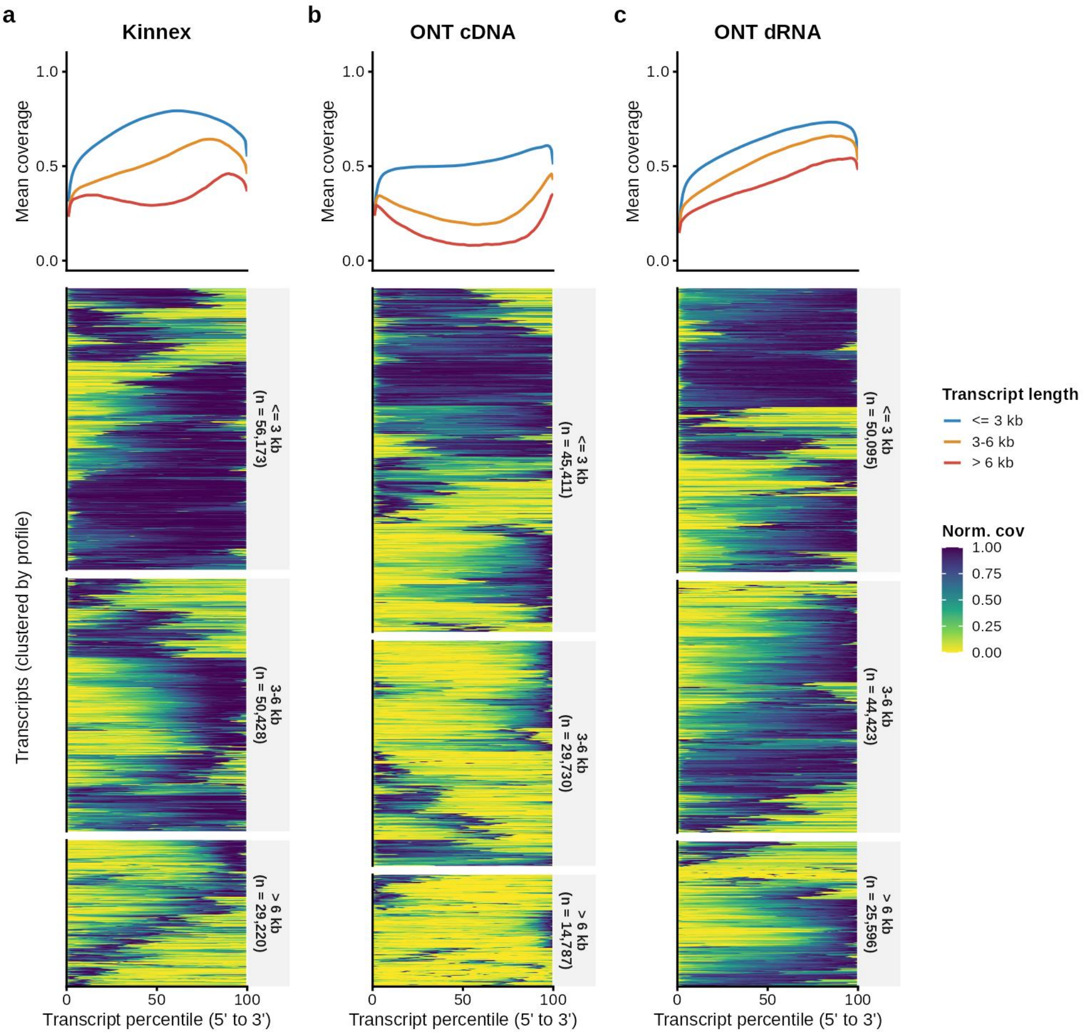


***Supplementary Fig. 3. Hierarchical clustering of individual transcript coverage profiles in the LongBench lung cancer cell line dataset***. Relative read coverage across transcript percentiles for individual transcripts clustered hierarchically across PacBio Kinnex, ONT cDNA, and ONT dRNA technologies, stratified by transcript length bins. Individual rows represent single transcript models.

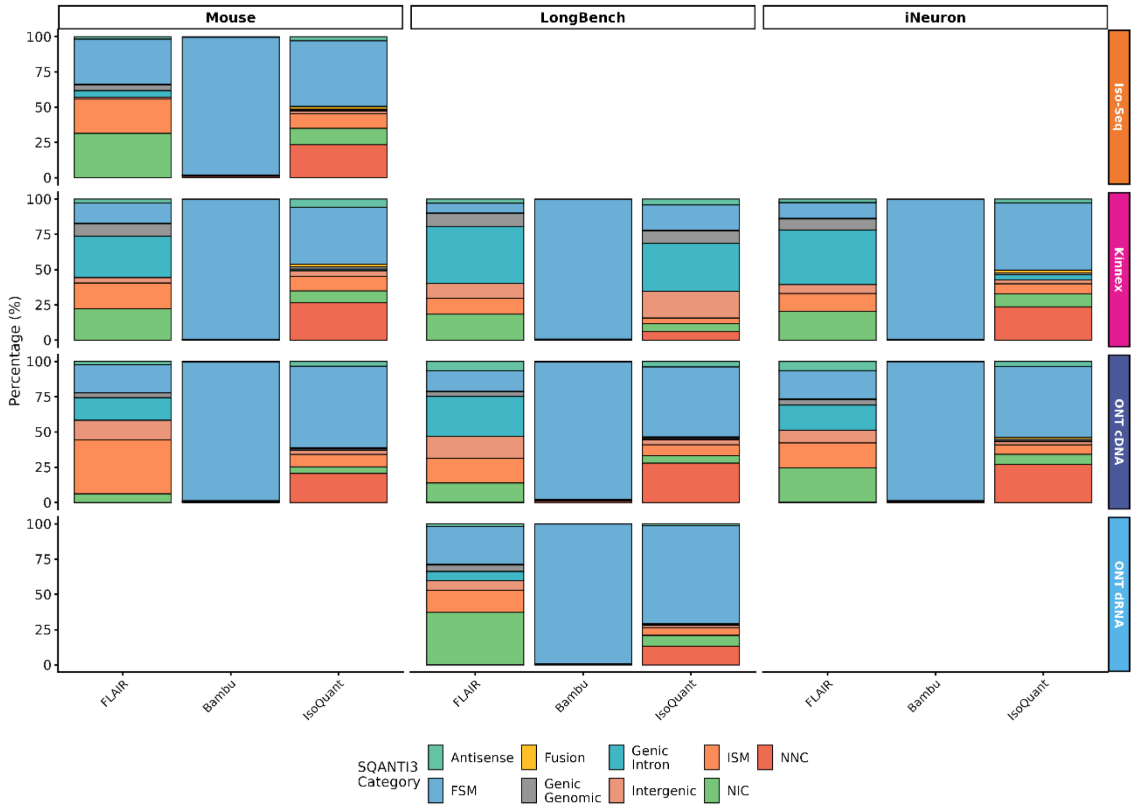


***Supplementary Fig. 4. SQANTI3 structural classification of unfiltered transcriptomes***. Distribution of transcript structural categories (including Full Splice Match [FSM], Incomplete Splice Match [ISM], Novel in Catalog [NIC], Novel Not in Catalog [NNC], and others) in the raw, unfiltered transcriptomes generated by bambu, FLAIR, and IsoQuant across technologies and datasets.

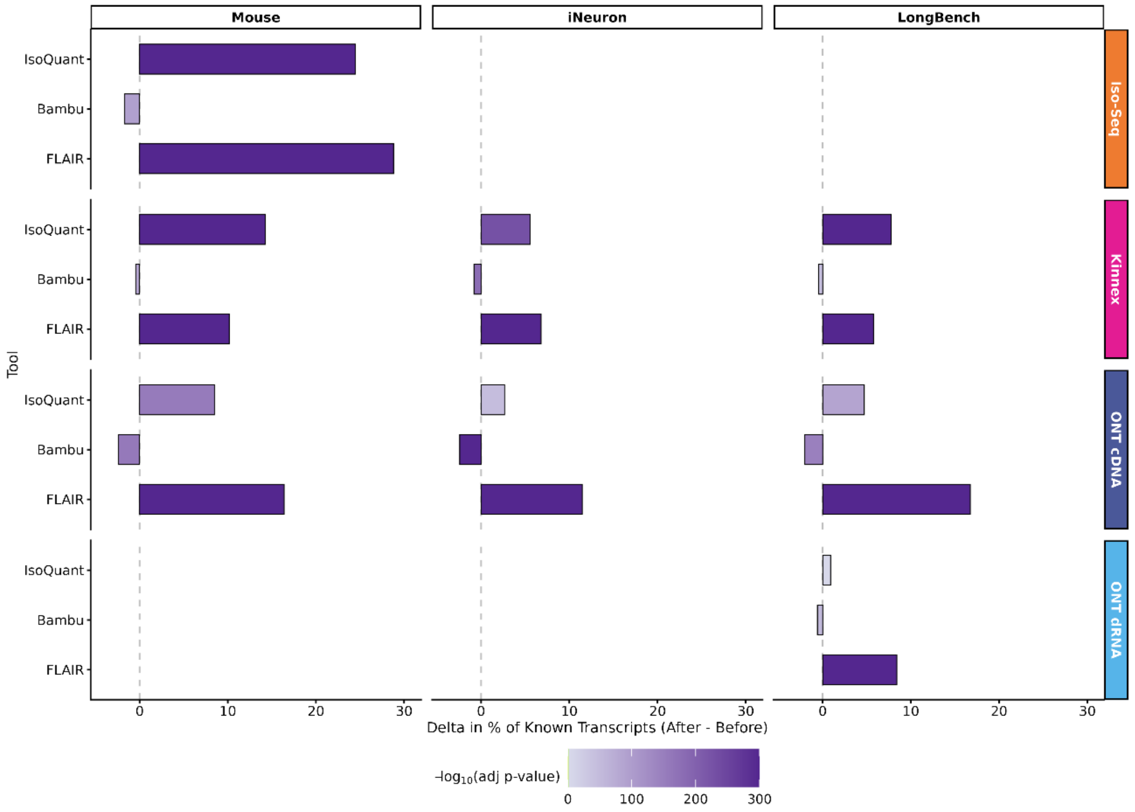


***Supplementary Fig. 5. Change in proportion of novel transcripts after abundance-based filtering***. Change across computational tools following expression-based filtering. Bar fill represents statistical significance level (-$\log_{10}$ p-value) from contingency table chi-squared tests comparing the proportion of novel transcripts before and after filtering.

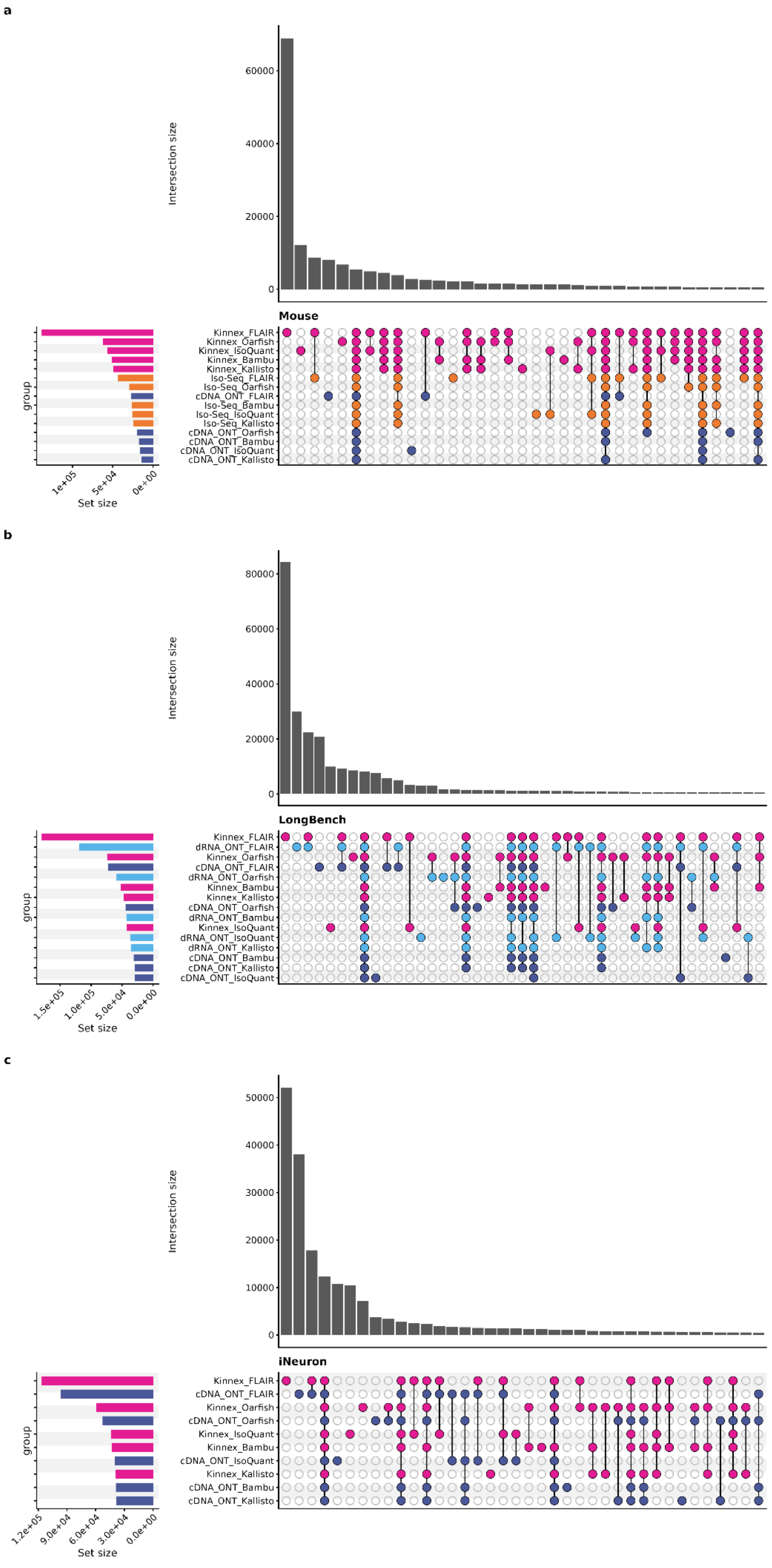
a
Intersection size
60000
40000
20000
0
Mouse
Kinnex_FLAIR
Kinnex_Oarfish
Kinnex_IsoQuant
Kinnex_Bambu
Kinnex_Kallisto
Iso-Seq_FLAIR
Iso-Seq_Oarfish
cDNA_ONT_FLAIR
Iso-Seq_Bambu
Iso-Seq_IsoQuant
Iso-Seq_Kallisto
cDNA_ONT_Oarfish
cDNA_ONT_Bambu
cDNA_ONT_IsoQuant
cDNA_ONT_Kallisto
group
1e+05
5e+04
0e+00
Set size
b
Intersection size
80000
60000
40000
20000
0
LongBench
Kinnex_FLAIR
dRNA_ONT_FLAIR
Kinnex_Oarfish
cDNA_ONT_FLAIR
dRNA_ONT_Oarfish
Kinnex_Bambu
Kinnex_Kallisto
cDNA_ONT_Oarfish
dRNA_ONT_Bambu
Kinnex_IsoQuant
dRNA_ONT_IsoQuant
dRNA_ONT_Kallisto
cDNA_ONT_Bambu
cDNA_ONT_Kallisto
cDNA_ONT_IsoQuant
group
1.5e+05
1.0e+05
5.0e+04
0.0e+00
Set size
c
Intersection size
50000
40000
30000
20000
10000
0
iNeuron
Kinnex_FLAIR
cDNA_ONT_FLAIR
Kinnex_Oarfish
cDNA_ONT_Oarfish
Kinnex_IsoQuant
Kinnex_Bambu
cDNA_ONT_IsoQuant
Kinnex_Kallisto
cDNA_ONT_Bambu
cDNA_ONT_Kallisto
group
1.2e+05
9.0e+04
6.0e+04
3.0e+04
0.0e+00
Set size

***Supplementary Fig. 6. Intersection of unique intron chains (UICs) across sequencing platforms and computational tools.*** Splicing model discovery consistency shown as intersections of unique intron chains (UICs) discovered by combinations of tools and sequencing technologies. Panels show: **a,** mouse tissue mix; **b,** human iNeuron; and **c,** LongBench cancer cell lines. Vertical bars represent intersection sizes, and horizontal bars represent the total number of unique intron chains discovered per tool-platform combination.

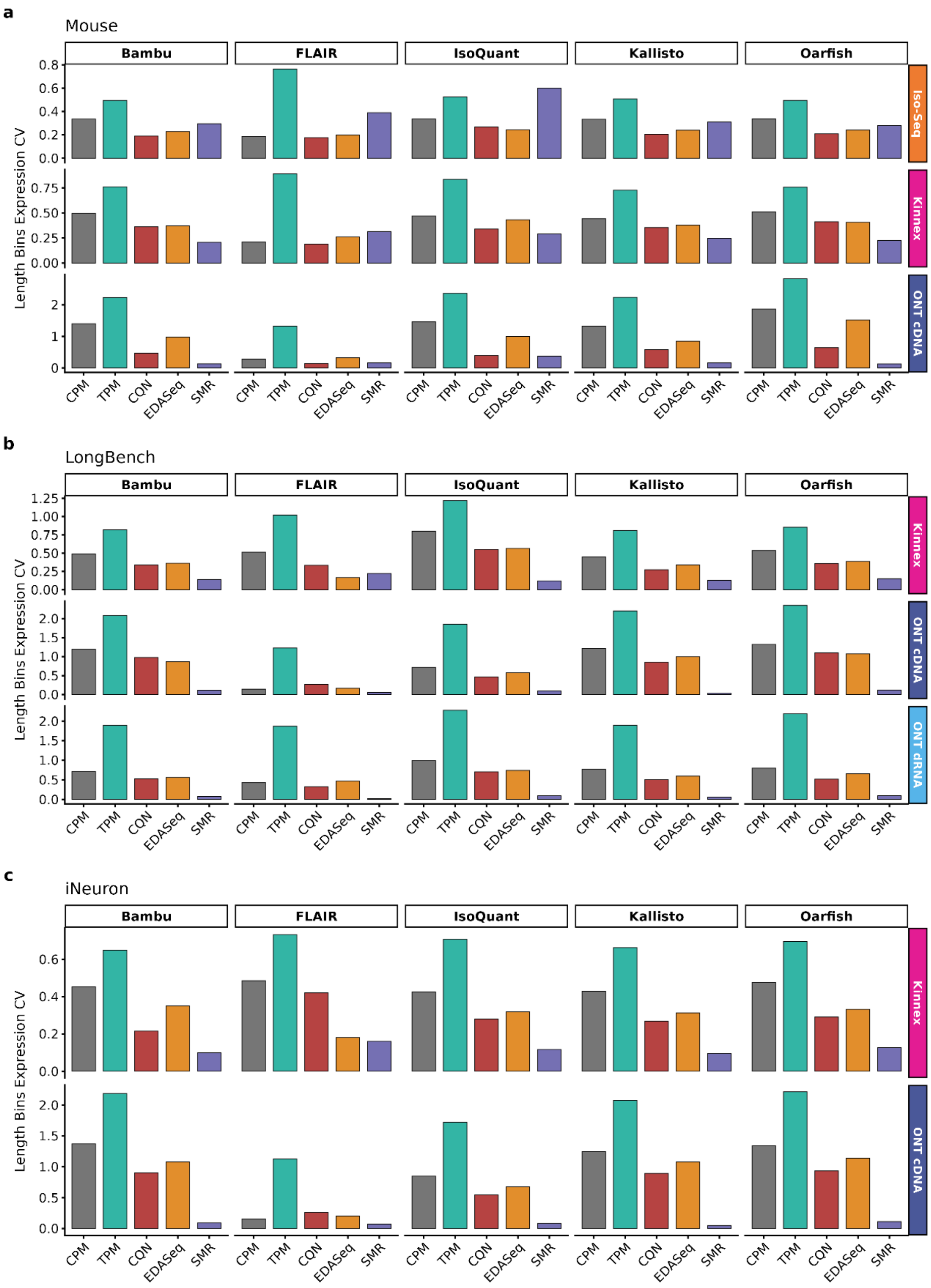


***Supplementary Fig. 7. Coefficient of Variation (CV) of mean expression across transcript length bins*** Coefficient of Variation of mean expression across transcript length bins for the assessed normalization methods. Rows represent sequencing technologies and columns represent upstream computational tools, for cohorts: **a,** mouse (Brain); **b,** LongBench (SCLC-A); and **c,** iNeurons (isoB11).

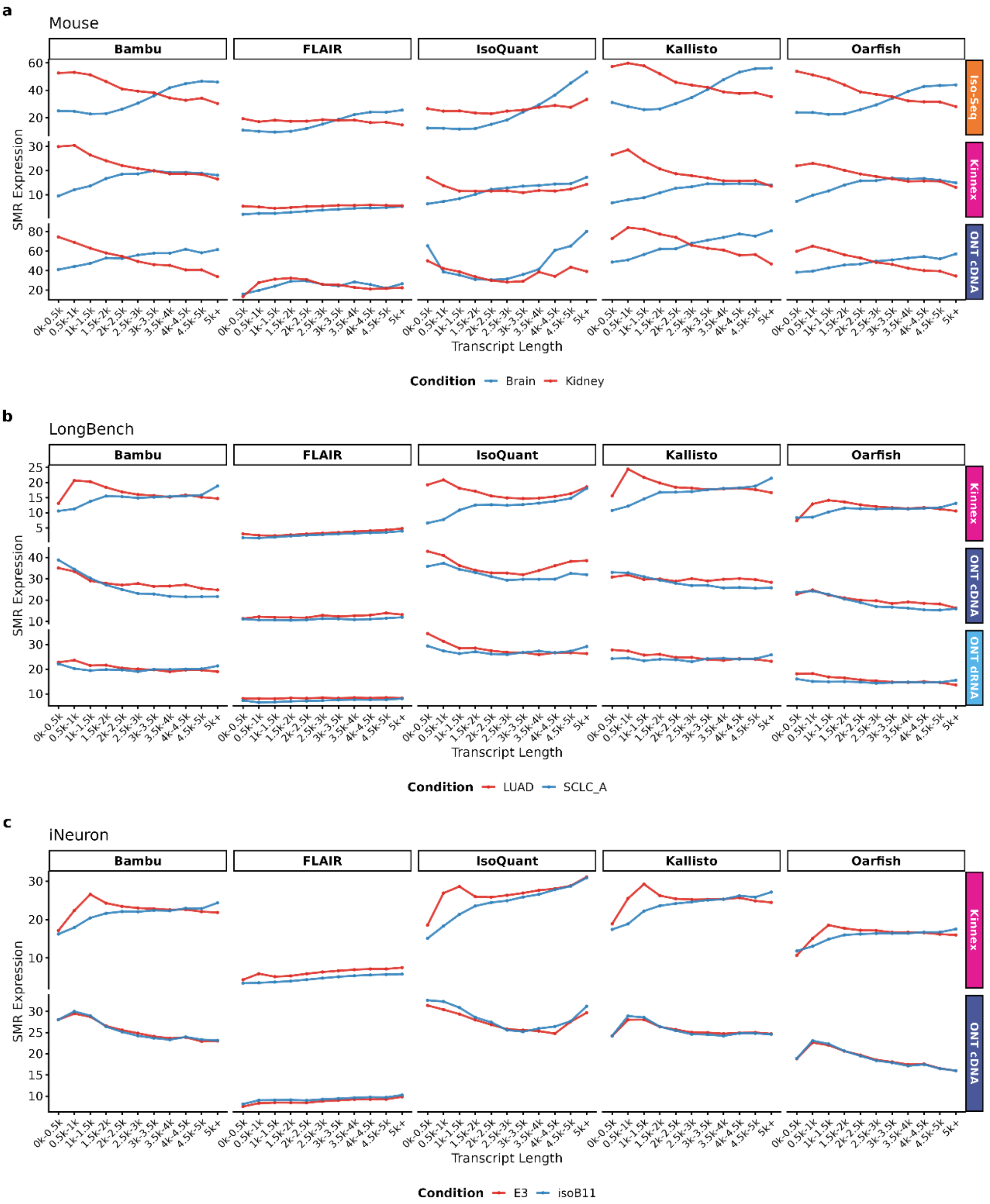


***Supplementary Fig. 8. Transcript length-expression profiles under Scaled Median Ratio (SMR) normalization.*** Trimmed mean of normalized expression across transcript length bins under SMR normalization, comparing different conditions in each dataset. Rows represent sequencing technologies and columns represent upstream computational tools, for cohorts: **a,** mouse; **b,** LongBench ; and **c,** iNeurons.

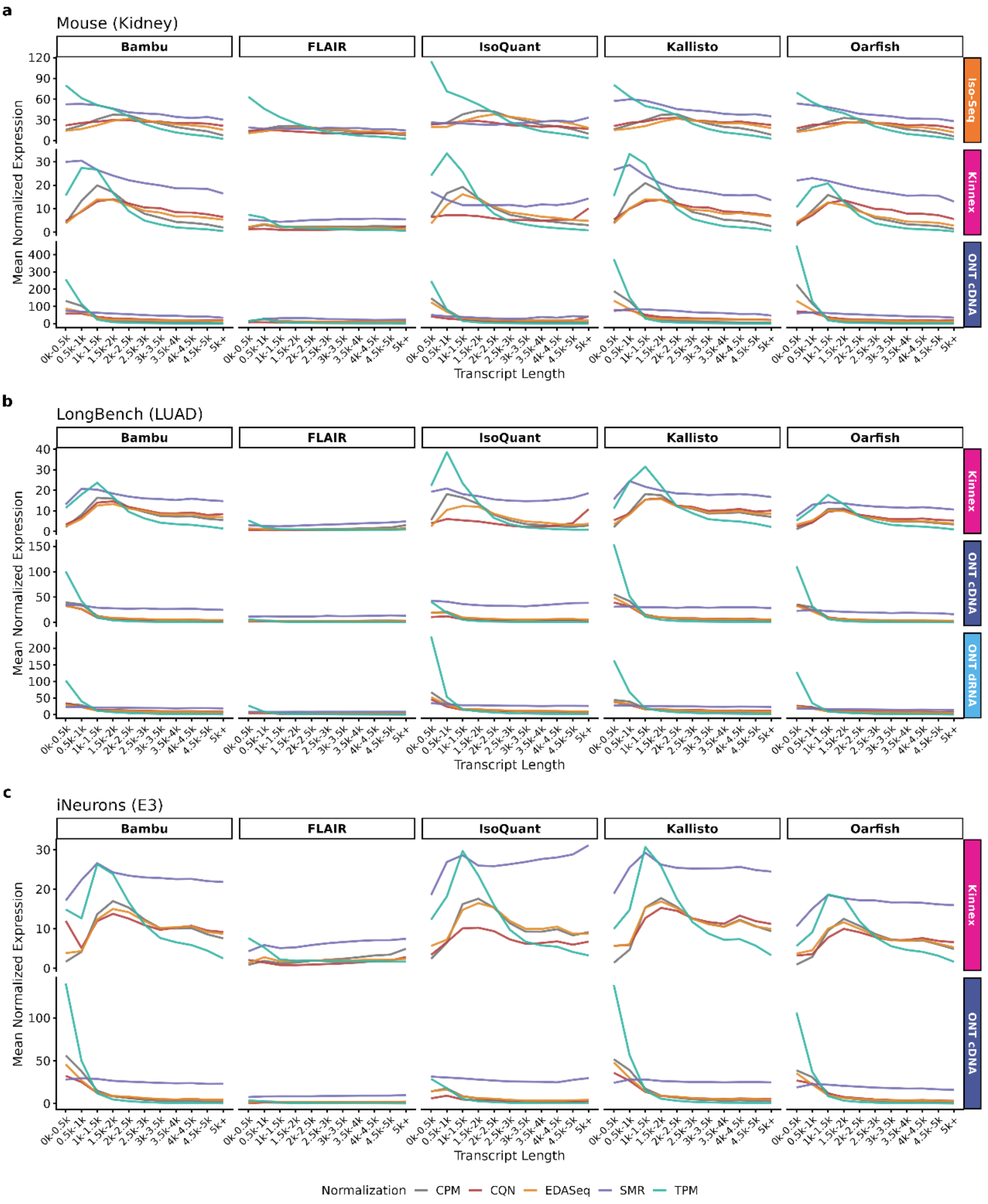


***Supplementary Fig. 9. Transcript length-expression relationship under different normalizations.*** Mean normalized expression across transcript length bins for five normalization methods: CPM, TPM, CQN, EDASeq, and Scaled Median Ratio (SMR). Panels represent different datasets: **a,** mouse Kidney; **b,** LongBench (LUAD); and **c,** human iNeuron (E3).

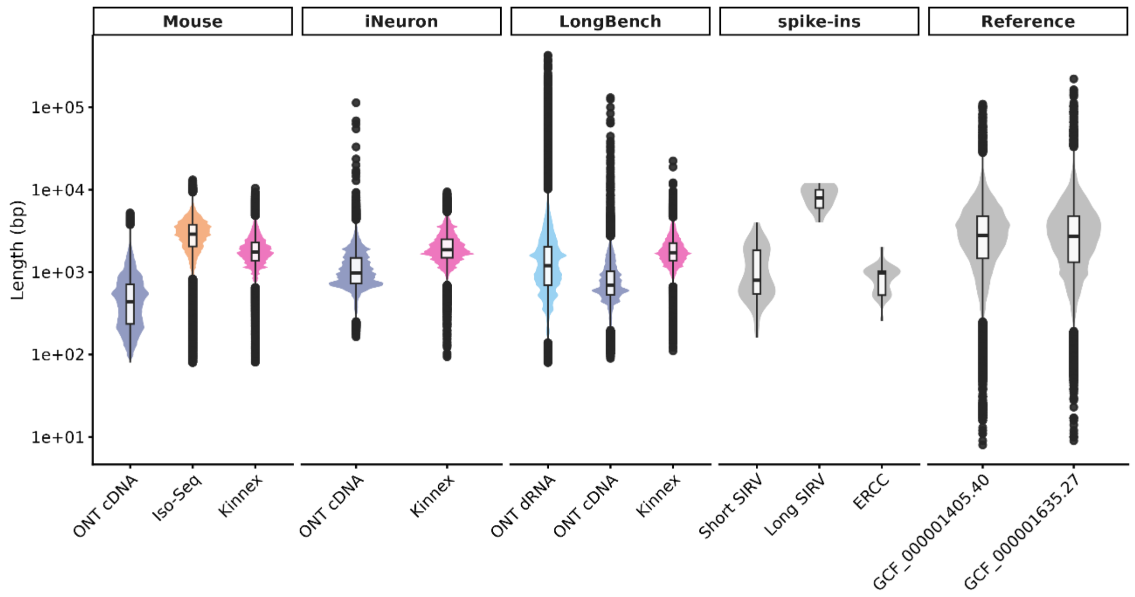


***Supplementary Fig. 10. Distribution of mapped read lengths across sequencing technologies and datasets.*** Read-length distributions for ONT dRNA, ONT cDNA, PacBio Iso-Seq and Kinnex libraries in the Mouse, iNeuron and LongBench cohorts, together with the reference length distributions of the SIRV and ERCC spike-in controls used for absolute quantification validation, and the reference annotation for human and mouse. For each library, 1 million mapped reads were randomly subsampled for plotting. Violin plots show the kernel density distribution of values; embedded box plots show the median, interquartile range and whiskers extending to the most extreme values within 1.5× the interquartile range.

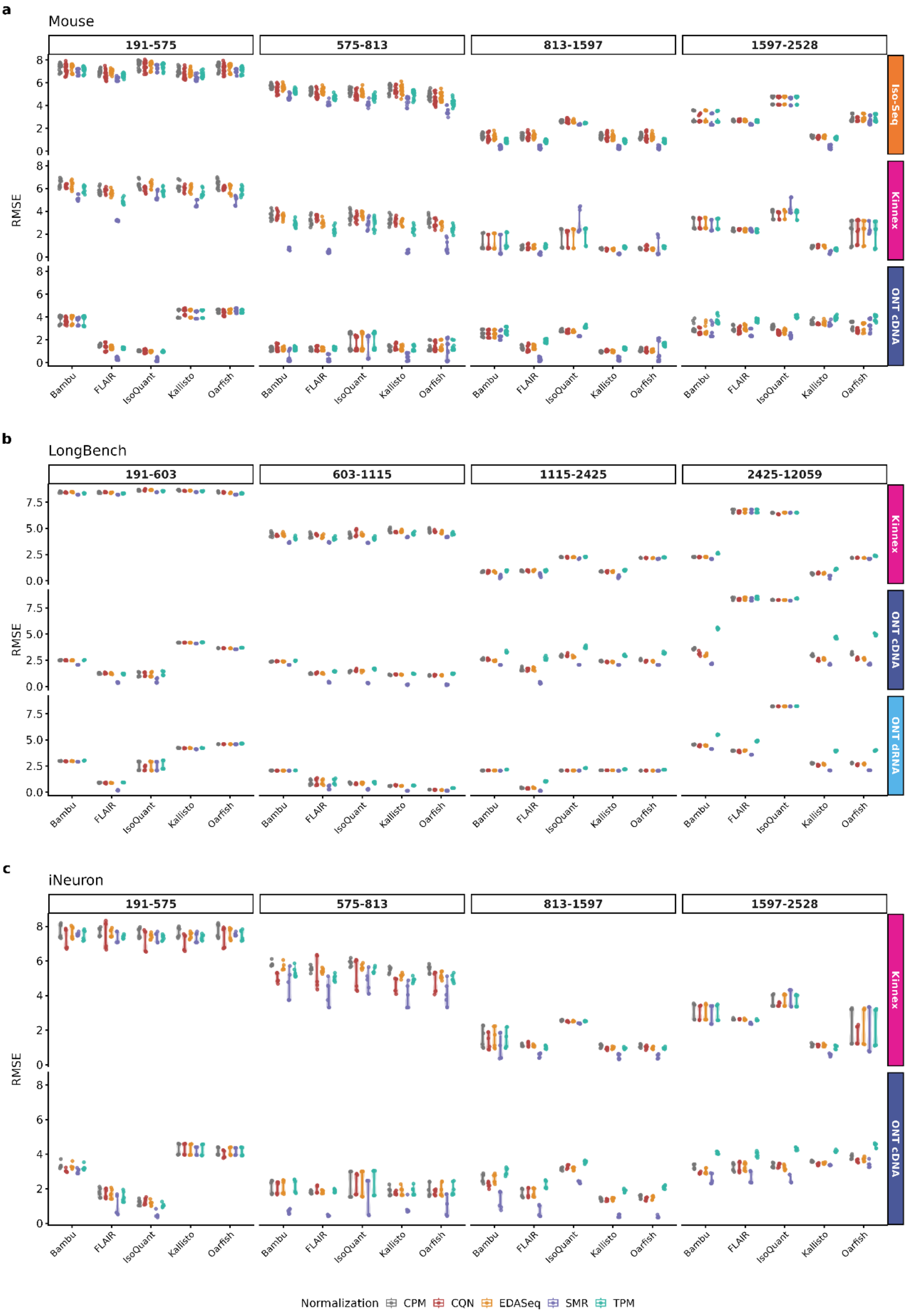


***Supplementary Fig. 11. Transcript-level absolute quantification error of SIRVs stratified by transcript length quantiles.*** Root-Mean-Square Error (RMSE) of SIRVs transcript quantification across upstream tools using different normalization methods. Panels show different datasets: **a**, mouse (n = 49 samples: 20 Iso-Seq, 9 Kinnex and 20 ONT cDNA); **b**, LongBench (n = 18: 6 Kinnex, 6 ONT cDNA and 6 ONT dRNA); **c**, human iNeuron (n = 12: 6 Kinnex and

6 ONT cDNA). Violin plots show the kernel density distribution of values; embedded box plots show the median, interquartile range and whiskers extending to the most extreme values within 1.5× the interquartile range.

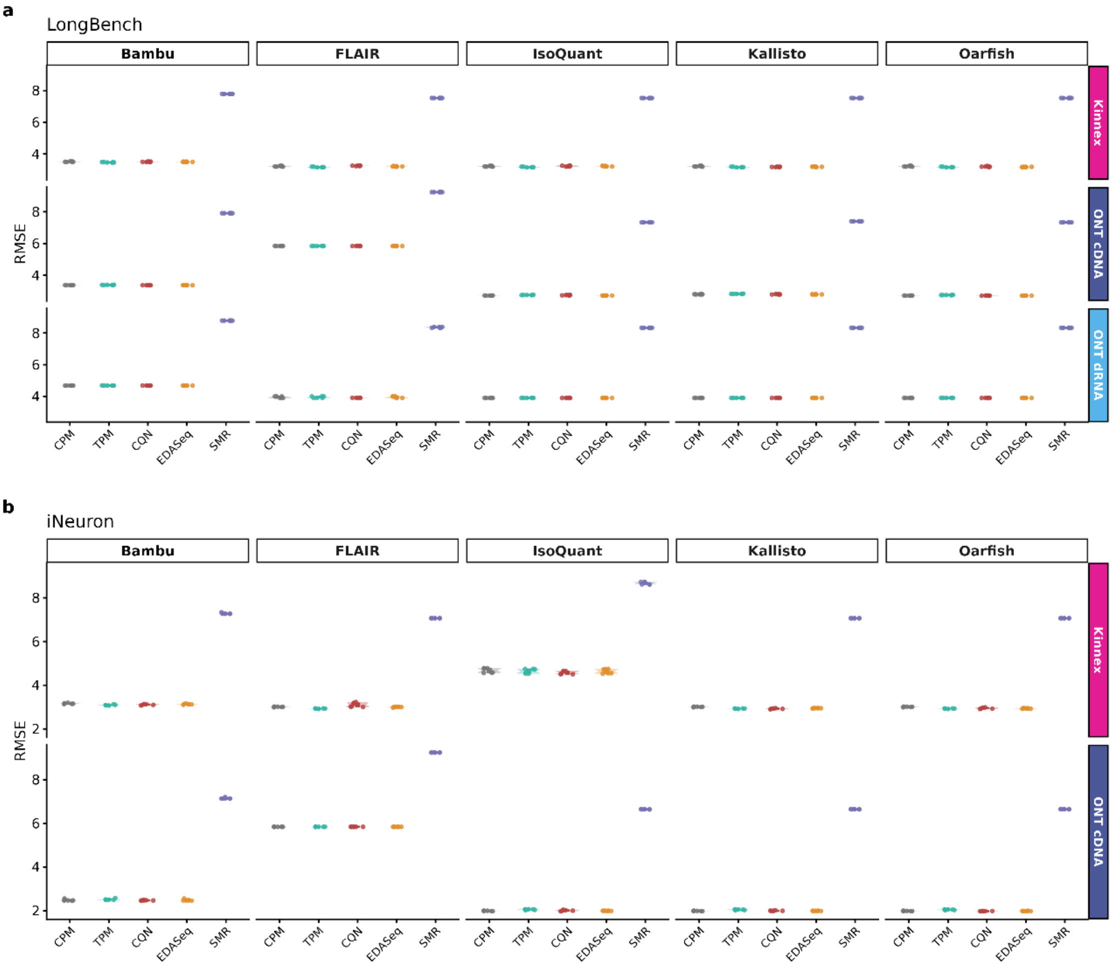


***Supplementary Fig. 12. Absolute quantification error of ERCC spike-ins.*** Root-Mean-Square Error (RMSE) of ERCC single-isoform spike-in controls across tools and normalization methods. Panels show different datasets **a**, LongBench (n = 18: 6 Kinnex, 6 ONT cDNA and 6 ONT dRNA); **b**, human iNeuron (n = 12: 6 Kinnex and 6 ONT cDNA). Violin plots show the kernel density distribution of values; embedded box plots show the median, interquartile range and whiskers extending to the most extreme values within 1.5× the interquartile range.

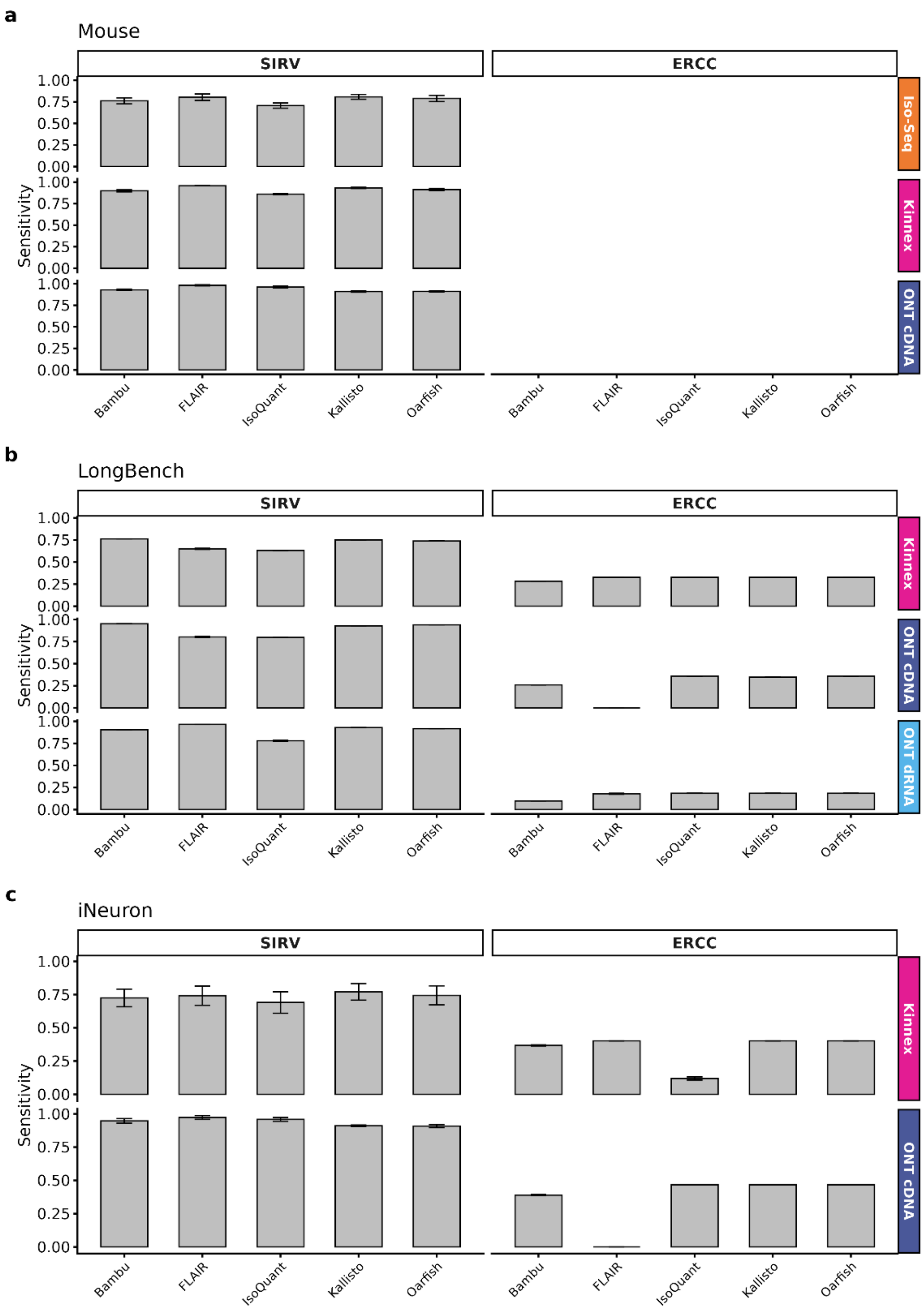


***Supplementary Fig. 13. Detection sensitivity of synthetic spike-ins.*** Sensitivity (fraction of spike-in transcripts detected) across computational tools and sequencing technologies, stratified by spike-in control type (SIRV or ERCC). Bars show the mean and error bars ± s.d. across samples. Panels show: **a**, mouse tissue mix (SIRV only); **b**, LongBench cancer cell lines (SIRV and ERCC); and **c**, human iNeuron (SIRV and ERCC). Sensitivity is computed over 69 SIRVs in **a** and **c**, 84 SIRVs (including long SIRVs) in **b**, and 92 ERCCs in **b** and **c**. $n = 20$ (Iso-Seq), 9 (Kinnex) and 20 (ONT cDNA) samples in **a**, and 6 samples per technology and spike-in type in **b** and **c**.

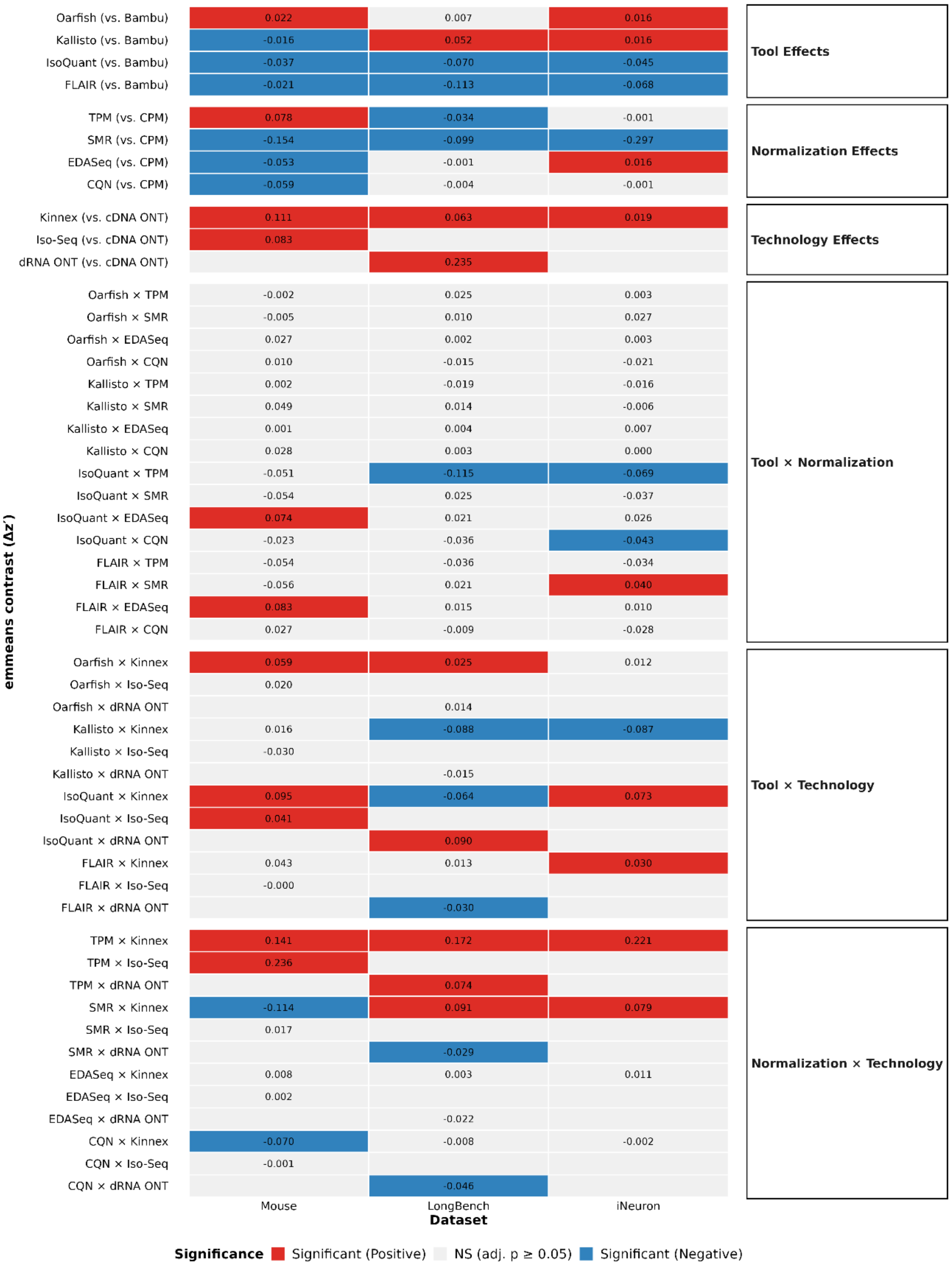


***Supplementary Fig. 14. Effects of tool, normalization and sequencing technology on short-read agreement estimated from the linear mixed model***. *Heatmap of estimated marginal mean (EMM) contrasts from linear mixed models fitted to Fisher z'-transformed transcript-level Spearman correlations against matched Illumina short-read data. Rows are grouped by term type. Main effects show each level relative to its reference (Bambu for tool, CPM for normalization, ONT cDNA for technology), averaged over the levels of the remaining factors. Two-way interaction terms show the difference between the effect of the first factor (vs. its reference) at the indicated level of the second factor and the same effect at the second factor's reference level. Cell values are contrast estimates (Δz'). Red and blue denote significant*

*positive and negative contrasts, respectively (Holm-adjusted $p < 0.05$ within each term type); grey denotes non-significant contrasts. Blank grey cells correspond to technologies not sequenced in that cohort.*

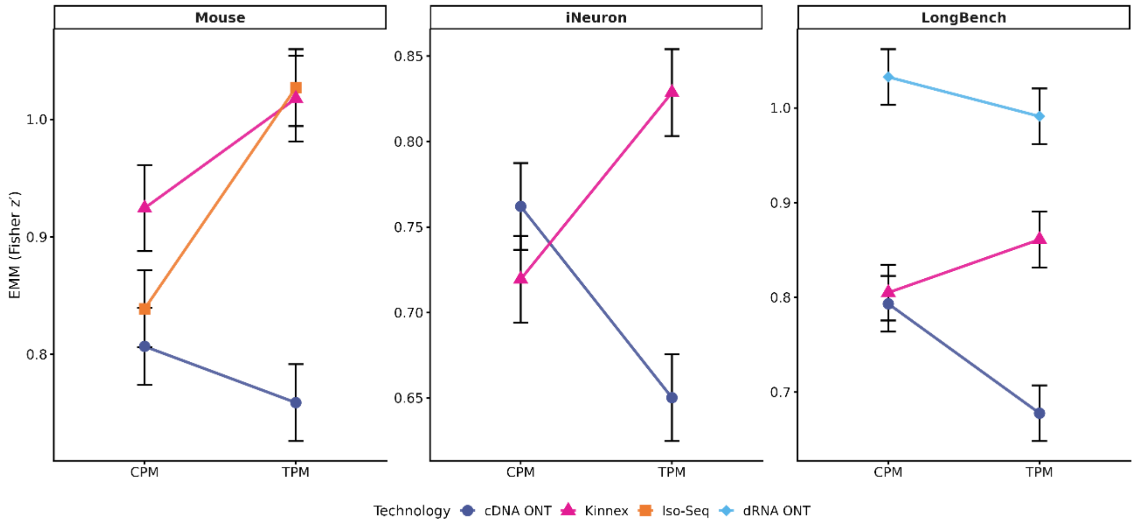


***Supplementary Fig. 15. Interaction between TPM normalization and sequencing technology.*** Predicted Spearman correlation coefficients (z-transformed) derived from mixed-effects models across normalizations and sequencing technologies. Points represent estimated marginal means (EMMs), and error bars represent 95% confidence intervals.

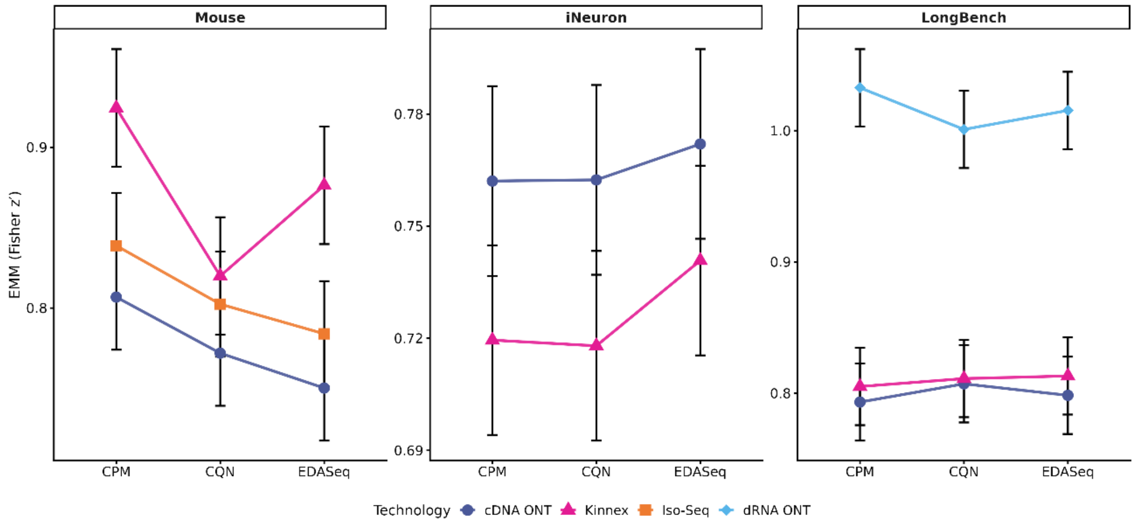


***Supplementary Fig. 16. Interaction between covariate-aware normalizations and sequencing technology.*** Predicted Spearman correlation coefficients (z-transformed) derived from mixed-effects models across normalizations and sequencing technologies. Points represent estimated marginal means (EMMs), and error bars represent 95% confidence intervals.

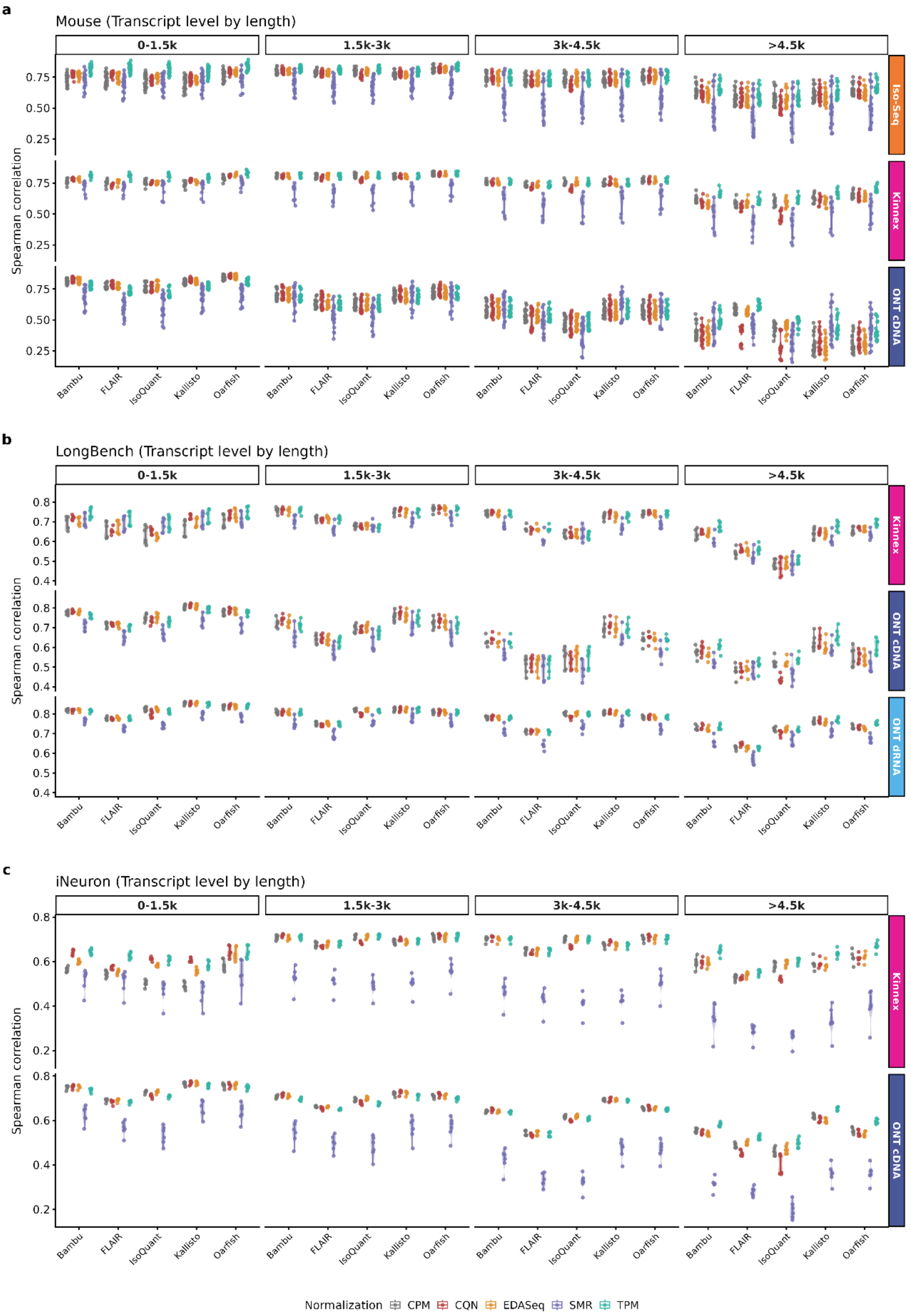


***Supplementary Fig. 17. Orthogonal validation correlations at the transcript level against matched Illumina short-read benchmarks stratified by transcript length quantiles.*** Spearman rank correlation coefficients between long-read expression estimates and matched Illumina short-read data across computational tools and normalization methods. Panels show different datasets: **a**, mouse (n = 49 samples: 20 Iso-Seq, 9 Kinnex and 20 ONT cDNA); **b**, LongBench (n

= 18: 6 Kinnex, 6 ONT cDNA and 6 ONT dRNA); **c**, human iNeuron (n = 12: 6 Kinnex and 6 ONT cDNA). Violin plots show the kernel density distribution of values; embedded box plots show the median, interquartile range and whiskers extending to the most extreme values within 1.5× the interquartile range.
