## Supplementary Tables 1-3 for "Systematic evaluation of transcript normalization methods and quantification pipelines across long-read sequencing platforms"

### Supplementary Table 1

**Table 1a. ANOVA (Type III)**

| Cohort | Term | Sum Sq | Mean Sq | Num df | Den df | F | p |
|---|---|---|---|---|---|---|---|
| Mouse | Normalization | 6.161 | 1.54 | 4 | 1163 | 277.6 | 1.52e-167 |
| Mouse | Platform | 2.433 | 1.217 | 2 | 1168 | 219.3 | 1.36e-81 |
| Mouse | Normalization × Platform | 2.72 | 0.34 | 8 | 1163 | 61.27 | 1.32e-83 |
| Mouse | Tool | 0.4312 | 0.1078 | 4 | 1163 | 19.43 | 1.72e-15 |
| Mouse | Tool × Normalization | 0.8339 | 0.05212 | 16 | 1163 | 9.39 | 2.67e-22 |
| Mouse | Tool × Platform | 0.2491 | 0.03114 | 8 | 1163 | 5.61 | 5.27e-07 |
| iNeuron | Normalization | 4.338 | 1.084 | 4 | 261 | 2062 | 4.3e-196 |
| iNeuron | Normalization × Platform | 0.5409 | 0.1352 | 4 | 261 | 257.1 | 3.04e-89 |
| iNeuron | Tool | 0.3478 | 0.08695 | 4 | 261 | 165.3 | 2.72e-70 |
| iNeuron | Tool × Platform | 0.2055 | 0.05137 | 4 | 261 | 97.68 | 1.09e-50 |
| iNeuron | Platform | 0.028 | 0.028 | 1 | 261 | 53.25 | 3.54e-12 |
| iNeuron | Tool × Normalization | 0.07875 | 0.004922 | 16 | 261 | 9.36 | 2.99e-18 |
| LongBench | Tool | 1.561 | 0.3901 | 4 | 392 | 550.1 | 2.61e-159 |
| LongBench | Normalization | 0.6462 | 0.1615 | 4 | 392 | 227.8 | 7.61e-101 |
| LongBench | Platform | 0.1177 | 0.05886 | 2 | 15 | 83 | 7.73e-09 |
| LongBench | Normalization × Platform | 0.4357 | 0.05446 | 8 | 392 | 76.8 | 1.64e-75 |
| LongBench | Tool × Platform | 0.4347 | 0.05434 | 8 | 392 | 76.63 | 2.13e-75 |
| LongBench | Tool × Normalization | 0.1974 | 0.01234 | 16 | 392 | 17.4 | 1.21e-36 |

**Table 1b. R2 (Nakagawa)**

| Cohort | Marginal R2 | Conditional R2 |
|---|---|---|
| Mouse | 0.522 | 0.723 |
| iNeuron | 0.944 | 0.973 |
| LongBench | 0.908 | 0.962 |

### Supplementary Table 2

**Table 2a. vs CPM (Dunnett)**

| Cohort | Platform | Contrast | Δz′ | SE | df | 95% CI lower | 95% CI upper | t | Adjusted p | Adjustment |
|---|---|---|---|---|---|---|---|---|---|---|
| Mouse | cDNA ONT | CQN - CPM | -0.0349 | 0.0105 | 1163 | -0.0608 | -0.009 | -3.31 | 0.00365 | Dunnett |
| Mouse | cDNA ONT | EDASeq - CPM | -0.0566 | 0.0105 | 1163 | -0.0825 | -0.0307 | -5.37 | 3.76e-07 | Dunnett |
| Mouse | cDNA ONT | SMR - CPM | -0.1211 | 0.0105 | 1163 | -0.147 | -0.0952 | -11.5 | < 1e-12 | Dunnett |
| Mouse | cDNA ONT | TPM - CPM | -0.0479 | 0.0105 | 1163 | -0.0738 | -0.022 | -4.55 | 2.39e-05 | Dunnett |
| Mouse | Iso-Seq | CQN - CPM | -0.0364 | 0.0105 | 1163 | -0.0622 | -0.0105 | -3.45 | 0.00222 | Dunnett |
| Mouse | Iso-Seq | EDASeq - CPM | -0.0548 | 0.0105 | 1163 | -0.0807 | -0.029 | -5.21 | 9.12e-07 | Dunnett |
| Mouse | Iso-Seq | SMR - CPM | -0.1045 | 0.0105 | 1163 | -0.1304 | -0.0787 | -9.92 | < 1e-12 | Dunnett |
| Mouse | Iso-Seq | TPM - CPM | 0.1882 | 0.0105 | 1163 | 0.1623 | 0.2141 | 17.87 | < 1e-12 | Dunnett |
| Mouse | Kinnex | CQN - CPM | -0.1046 | 0.0157 | 1163 | -0.1431 | -0.066 | -6.66 | 1.7e-10 | Dunnett |
| Mouse | Kinnex | EDASeq - CPM | -0.0482 | 0.0157 | 1163 | -0.0867 | -0.0096 | -3.07 | 0.00834 | Dunnett |
| Mouse | Kinnex | SMR - CPM | -0.2354 | 0.0157 | 1163 | -0.274 | -0.1968 | -14.99 | < 1e-12 | Dunnett |
| Mouse | Kinnex | TPM - CPM | 0.0932 | 0.0157 | 1163 | 0.0547 | 0.1318 | 5.94 | 1.52e-08 | Dunnett |
| iNeuron | cDNA ONT | CQN - CPM | 3.000e-04 | 0.0059 | 261 | -0.0143 | 0.0149 | 0.05 | 1 | Dunnett |
| iNeuron | cDNA ONT | EDASeq - CPM | 0.0099 | 0.0059 | 261 | -0.0047 | 0.0245 | 1.67 | 0.28 | Dunnett |
| iNeuron | cDNA ONT | SMR - CPM | -0.3362 | 0.0059 | 261 | -0.3509 | -0.3216 | -56.78 | < 1e-12 | Dunnett |
| iNeuron | cDNA ONT | TPM - CPM | -0.1119 | 0.0059 | 261 | -0.1265 | -0.0973 | -18.9 | < 1e-12 | Dunnett |
| iNeuron | Kinnex | CQN - CPM | -0.0015 | 0.0059 | 261 | -0.0161 | 0.0131 | -0.26 | 0.986 | Dunnett |
| iNeuron | Kinnex | EDASeq - CPM | 0.0213 | 0.0059 | 261 | 0.0067 | 0.0359 | 3.6 | 0.00148 | Dunnett |
| iNeuron | Kinnex | SMR - CPM | -0.2573 | 0.0059 | 261 | -0.2719 | -0.2427 | -43.46 | < 1e-12 | Dunnett |
| iNeuron | Kinnex | TPM - CPM | 0.1092 | 0.0059 | 261 | 0.0945 | 0.1238 | 18.43 | < 1e-12 | Dunnett |
| LongBench | cDNA ONT | CQN - CPM | 0.0139 | 0.0069 | 392 | -0.003 | 0.0308 | 2.02 | 0.143 | Dunnett |
| LongBench | cDNA ONT | EDASeq - CPM | 0.0051 | 0.0069 | 392 | -0.0118 | 0.0221 | 0.75 | 0.829 | Dunnett |
| LongBench | cDNA ONT | SMR - CPM | -0.1197 | 0.0069 | 392 | -0.1366 | -0.1028 | -17.41 | < 1e-12 | Dunnett |
| LongBench | cDNA ONT | TPM - CPM | -0.1155 | 0.0069 | 392 | -0.1325 | -0.0986 | -16.8 | < 1e-12 | Dunnett |
| LongBench | Kinnex | CQN - CPM | 0.006 | 0.0069 | 392 | -0.0109 | 0.023 | 0.88 | 0.758 | Dunnett |
| LongBench | Kinnex | EDASeq - CPM | 0.008 | 0.0069 | 392 | -0.0089 | 0.025 | 1.17 | 0.576 | Dunnett |
| LongBench | Kinnex | SMR - CPM | -0.029 | 0.0069 | 392 | -0.0459 | -0.012 | -4.21 | 0.000124 | Dunnett |
| LongBench | Kinnex | TPM - CPM | 0.0561 | 0.0069 | 392 | 0.0391 | 0.073 | 8.15 | < 1e-12 | Dunnett |
| LongBench | dRNA ONT | CQN - CPM | -0.0317 | 0.0069 | 392 | -0.0487 | -0.0148 | -4.62 | 2.09e-05 | Dunnett |
| LongBench | dRNA ONT | EDASeq - CPM | -0.0173 | 0.0069 | 392 | -0.0342 | -3.000e-04 | -2.51 | 0.044 | Dunnett |
| LongBench | dRNA ONT | SMR - CPM | -0.1491 | 0.0069 | 392 | -0.166 | -0.1321 | -21.68 | < 1e-12 | Dunnett |
| LongBench | dRNA ONT | TPM - CPM | -0.0415 | 0.0069 | 392 | -0.0585 | -0.0246 | -6.04 | 1.42e-08 | Dunnett |

**Table 2b. Pairwise (Tukey)**

| Cohort | Platform | Contrast | Δz′ | SE | df | 95% CI lower | 95% CI upper | t | Adjusted p | Adjustment |
|---|---|---|---|---|---|---|---|---|---|---|
| Mouse | cDNA ONT | CPM - CQN | 0.0349 | 0.0105 | 1163 | 0.0061 | 0.0637 | 3.31 | 0.00846 | Tukey |
| Mouse | cDNA ONT | CPM - EDASeq | 0.0566 | 0.0105 | 1163 | 0.0278 | 0.0854 | 5.37 | 9.38e-07 | Tukey |
| Mouse | cDNA ONT | CPM - SMR | 0.1211 | 0.0105 | 1163 | 0.0923 | 0.1499 | 11.5 | < 1e-12 | Tukey |
| Mouse | cDNA ONT | CPM - TPM | 0.0479 | 0.0105 | 1163 | 0.0191 | 0.0767 | 4.55 | 5.89e-05 | Tukey |
| Mouse | cDNA ONT | CQN - EDASeq | 0.0217 | 0.0105 | 1163 | -0.0071 | 0.0505 | 2.06 | 0.239 | Tukey |
| Mouse | cDNA ONT | CQN - SMR | 0.0862 | 0.0105 | 1163 | 0.0574 | 0.115 | 8.19 | < 1e-12 | Tukey |
| Mouse | cDNA ONT | CQN - TPM | 0.013 | 0.0105 | 1163 | -0.0158 | 0.0418 | 1.23 | 0.731 | Tukey |
| Mouse | cDNA ONT | EDASeq - SMR | 0.0645 | 0.0105 | 1163 | 0.0358 | 0.0933 | 6.13 | 1.23e-08 | Tukey |

| Cohort | Platform | Contrast | Δz′ | SE | df | 95% CI lower | 95% CI upper | t | Adjusted p | Adjustment |
|---|---|---|---|---|---|---|---|---|---|---|
| Mouse | cDNA ONT | EDASeq - TPM | -0.0087 | 0.0105 | 1163 | -0.0375 | 0.0201 | -0.82 | 0.923 | Tukey |
| Mouse | cDNA ONT | SMR - TPM | -0.0732 | 0.0105 | 1163 | -0.102 | -0.0444 | -6.95 | 6.11e-11 | Tukey |
| Mouse | Iso-Seq | CPM - CQN | 0.0364 | 0.0105 | 1163 | 0.0076 | 0.0651 | 3.45 | 0.0052 | Tukey |
| Mouse | Iso-Seq | CPM - EDASeq | 0.0548 | 0.0105 | 1163 | 0.0261 | 0.0836 | 5.21 | 2.27e-06 | Tukey |
| Mouse | Iso-Seq | CPM - SMR | 0.1045 | 0.0105 | 1163 | 0.0758 | 0.1333 | 9.92 | < 1e-12 | Tukey |
| Mouse | Iso-Seq | CPM - TPM | -0.1882 | 0.0105 | 1163 | -0.217 | -0.1594 | -17.87 | < 1e-12 | Tukey |
| Mouse | Iso-Seq | CQN - EDASeq | 0.0185 | 0.0105 | 1163 | -0.0103 | 0.0472 | 1.75 | 0.402 | Tukey |
| Mouse | Iso-Seq | CQN - SMR | 0.0682 | 0.0105 | 1163 | 0.0394 | 0.0969 | 6.47 | 1.43e-09 | Tukey |
| Mouse | Iso-Seq | CQN - TPM | -0.2246 | 0.0105 | 1163 | -0.2534 | -0.1958 | -21.32 | < 1e-12 | Tukey |
| Mouse | Iso-Seq | EDASeq - SMR | 0.0497 | 0.0105 | 1163 | 0.0209 | 0.0785 | 4.72 | 2.63e-05 | Tukey |
| Mouse | Iso-Seq | EDASeq - TPM | -0.2431 | 0.0105 | 1163 | -0.2718 | -0.2143 | -23.07 | < 1e-12 | Tukey |
| Mouse | Iso-Seq | SMR - TPM | -0.2928 | 0.0105 | 1163 | -0.3215 | -0.264 | -27.79 | < 1e-12 | Tukey |
| Mouse | Kinnex | CPM - CQN | 0.1046 | 0.0157 | 1163 | 0.0617 | 0.1475 | 6.66 | 4.25e-10 | Tukey |
| Mouse | Kinnex | CPM - EDASeq | 0.0482 | 0.0157 | 1163 | 0.0053 | 0.0911 | 3.07 | 0.0188 | Tukey |
| Mouse | Kinnex | CPM - SMR | 0.2354 | 0.0157 | 1163 | 0.1925 | 0.2783 | 14.99 | < 1e-12 | Tukey |
| Mouse | Kinnex | CPM - TPM | -0.0932 | 0.0157 | 1163 | -0.1361 | -0.0503 | -5.94 | 3.8e-08 | Tukey |
| Mouse | Kinnex | CQN - EDASeq | -0.0564 | 0.0157 | 1163 | -0.0993 | -0.0135 | -3.59 | 0.00314 | Tukey |
| Mouse | Kinnex | CQN - SMR | 0.1308 | 0.0157 | 1163 | 0.0879 | 0.1737 | 8.33 | < 1e-12 | Tukey |
| Mouse | Kinnex | CQN - TPM | -0.1978 | 0.0157 | 1163 | -0.2407 | -0.1549 | -12.6 | < 1e-12 | Tukey |
| Mouse | Kinnex | EDASeq - SMR | 0.1872 | 0.0157 | 1163 | 0.1443 | 0.2301 | 11.92 | < 1e-12 | Tukey |
| Mouse | Kinnex | EDASeq - TPM | -0.1414 | 0.0157 | 1163 | -0.1843 | -0.0985 | -9 | < 1e-12 | Tukey |
| Mouse | Kinnex | SMR - TPM | -0.3286 | 0.0157 | 1163 | -0.3715 | -0.2857 | -20.93 | < 1e-12 | Tukey |
| iNeuron | cDNA ONT | CPM - CQN | -3.000e-04 | 0.0059 | 261 | -0.0166 | 0.016 | -0.05 | 1 | Tukey |
| iNeuron | cDNA ONT | CPM - EDASeq | -0.0099 | 0.0059 | 261 | -0.0262 | 0.0064 | -1.67 | 0.452 | Tukey |
| iNeuron | cDNA ONT | CPM - SMR | 0.3362 | 0.0059 | 261 | 0.32 | 0.3525 | 56.78 | < 1e-12 | Tukey |
| iNeuron | cDNA ONT | CPM - TPM | 0.1119 | 0.0059 | 261 | 0.0956 | 0.1282 | 18.9 | < 1e-12 | Tukey |
| iNeuron | cDNA ONT | CQN - EDASeq | -0.0096 | 0.0059 | 261 | -0.0259 | 0.0067 | -1.62 | 0.485 | Tukey |
| iNeuron | cDNA ONT | CQN - SMR | 0.3366 | 0.0059 | 261 | 0.3203 | 0.3528 | 56.84 | < 1e-12 | Tukey |
| iNeuron | cDNA ONT | CQN - TPM | 0.1122 | 0.0059 | 261 | 0.0959 | 0.1285 | 18.95 | < 1e-12 | Tukey |
| iNeuron | cDNA ONT | EDASeq - SMR | 0.3462 | 0.0059 | 261 | 0.3299 | 0.3624 | 58.46 | < 1e-12 | Tukey |
| iNeuron | cDNA ONT | EDASeq - TPM | 0.1218 | 0.0059 | 261 | 0.1055 | 0.1381 | 20.57 | < 1e-12 | Tukey |
| iNeuron | cDNA ONT | SMR - TPM | -0.2244 | 0.0059 | 261 | -0.2406 | -0.2081 | -37.89 | < 1e-12 | Tukey |
| iNeuron | Kinnex | CPM - CQN | 0.0015 | 0.0059 | 261 | -0.0147 | 0.0178 | 0.26 | 0.999 | Tukey |
| iNeuron | Kinnex | CPM - EDASeq | -0.0213 | 0.0059 | 261 | -0.0376 | -0.005 | -3.6 | 0.00349 | Tukey |
| iNeuron | Kinnex | CPM - SMR | 0.2573 | 0.0059 | 261 | 0.2411 | 0.2736 | 43.46 | < 1e-12 | Tukey |
| iNeuron | Kinnex | CPM - TPM | -0.1092 | 0.0059 | 261 | -0.1254 | -0.0929 | -18.43 | < 1e-12 | Tukey |
| iNeuron | Kinnex | CQN - EDASeq | -0.0228 | 0.0059 | 261 | -0.0391 | -0.0066 | -3.86 | 0.00136 | Tukey |
| iNeuron | Kinnex | CQN - SMR | 0.2558 | 0.0059 | 261 | 0.2395 | 0.2721 | 43.2 | < 1e-12 | Tukey |
| iNeuron | Kinnex | CQN - TPM | -0.1107 | 0.0059 | 261 | -0.127 | -0.0944 | -18.69 | < 1e-12 | Tukey |
| iNeuron | Kinnex | EDASeq - SMR | 0.2786 | 0.0059 | 261 | 0.2624 | 0.2949 | 47.06 | < 1e-12 | Tukey |
| iNeuron | Kinnex | EDASeq - TPM | -0.0878 | 0.0059 | 261 | -0.1041 | -0.0716 | -14.84 | < 1e-12 | Tukey |
| iNeuron | Kinnex | SMR - TPM | -0.3665 | 0.0059 | 261 | -0.3828 | -0.3502 | -61.89 | < 1e-12 | Tukey |
| LongBench | cDNA ONT | CPM - CQN | -0.0139 | 0.0069 | 392 | -0.0327 | 0.005 | -2.02 | 0.258 | Tukey |
| LongBench | cDNA ONT | CPM - EDASeq | -0.0051 | 0.0069 | 392 | -0.024 | 0.0137 | -0.75 | 0.946 | Tukey |

| Cohort | Platform | Contrast | Δz′ | SE | df | 95% CI lower | 95% CI upper | t | Adjusted p | Adjustment |
|---|---|---|---|---|---|---|---|---|---|---|
| LongBench | cDNA ONT | CPM - SMR | 0.1197 | 0.0069 | 392 | 0.1009 | 0.1386 | 17.41 | < 1e-12 | Tukey |
| LongBench | cDNA ONT | CPM - TPM | 0.1155 | 0.0069 | 392 | 0.0967 | 0.1344 | 16.8 | < 1e-12 | Tukey |
| LongBench | cDNA ONT | CQN - EDASeq | 0.0088 | 0.0069 | 392 | -0.0101 | 0.0276 | 1.27 | 0.707 | Tukey |
| LongBench | cDNA ONT | CQN - SMR | 0.1336 | 0.0069 | 392 | 0.1148 | 0.1524 | 19.43 | < 1e-12 | Tukey |
| LongBench | cDNA ONT | CQN - TPM | 0.1294 | 0.0069 | 392 | 0.1106 | 0.1483 | 18.82 | < 1e-12 | Tukey |
| LongBench | cDNA ONT | EDASeq - SMR | 0.1248 | 0.0069 | 392 | 0.106 | 0.1437 | 18.16 | < 1e-12 | Tukey |
| LongBench | cDNA ONT | EDASeq - TPM | 0.1207 | 0.0069 | 392 | 0.1018 | 0.1395 | 17.55 | < 1e-12 | Tukey |
| LongBench | cDNA ONT | SMR - TPM | -0.0042 | 0.0069 | 392 | -0.023 | 0.0147 | -0.61 | 0.974 | Tukey |
| LongBench | Kinnex | CPM - CQN | -0.006 | 0.0069 | 392 | -0.0249 | 0.0128 | -0.88 | 0.905 | Tukey |
| LongBench | Kinnex | CPM - EDASeq | -0.008 | 0.0069 | 392 | -0.0269 | 0.0108 | -1.17 | 0.77 | Tukey |
| LongBench | Kinnex | CPM - SMR | 0.029 | 0.0069 | 392 | 0.0101 | 0.0478 | 4.21 | 0.000302 | Tukey |
| LongBench | Kinnex | CPM - TPM | -0.0561 | 0.0069 | 392 | -0.0749 | -0.0372 | -8.15 | < 1e-12 | Tukey |
| LongBench | Kinnex | CQN - EDASeq | -0.002 | 0.0069 | 392 | -0.0208 | 0.0168 | -0.29 | 0.998 | Tukey |
| LongBench | Kinnex | CQN - SMR | 0.035 | 0.0069 | 392 | 0.0161 | 0.0538 | 5.09 | 5.54e-06 | Tukey |
| LongBench | Kinnex | CQN - TPM | -0.05 | 0.0069 | 392 | -0.0689 | -0.0312 | -7.28 | 2.08e-12 | Tukey |
| LongBench | Kinnex | EDASeq - SMR | 0.037 | 0.0069 | 392 | 0.0182 | 0.0558 | 5.38 | 1.27e-06 | Tukey |
| LongBench | Kinnex | EDASeq - TPM | -0.048 | 0.0069 | 392 | -0.0669 | -0.0292 | -6.99 | 1.05e-10 | Tukey |
| LongBench | Kinnex | SMR - TPM | -0.085 | 0.0069 | 392 | -0.1039 | -0.0662 | -12.37 | < 1e-12 | Tukey |
| LongBench | dRNA ONT | CPM - CQN | 0.0317 | 0.0069 | 392 | 0.0129 | 0.0506 | 4.62 | 5.17e-05 | Tukey |
| LongBench | dRNA ONT | CPM - EDASeq | 0.0173 | 0.0069 | 392 | -0.0016 | 0.0361 | 2.51 | 0.0901 | Tukey |
| LongBench | dRNA ONT | CPM - SMR | 0.1491 | 0.0069 | 392 | 0.1302 | 0.1679 | 21.68 | < 1e-12 | Tukey |
| LongBench | dRNA ONT | CPM - TPM | 0.0415 | 0.0069 | 392 | 0.0227 | 0.0604 | 6.04 | 3.55e-08 | Tukey |
| LongBench | dRNA ONT | CQN - EDASeq | -0.0145 | 0.0069 | 392 | -0.0333 | 0.0044 | -2.11 | 0.22 | Tukey |
| LongBench | dRNA ONT | CQN - SMR | 0.1173 | 0.0069 | 392 | 0.0985 | 0.1362 | 17.06 | < 1e-12 | Tukey |
| LongBench | dRNA ONT | CQN - TPM | 0.0098 | 0.0069 | 392 | -0.0091 | 0.0286 | 1.42 | 0.613 | Tukey |
| LongBench | dRNA ONT | EDASeq - SMR | 0.1318 | 0.0069 | 392 | 0.113 | 0.1506 | 19.17 | < 1e-12 | Tukey |
| LongBench | dRNA ONT | EDASeq - TPM | 0.0243 | 0.0069 | 392 | 0.0054 | 0.0431 | 3.53 | 0.00423 | Tukey |
| LongBench | dRNA ONT | SMR - TPM | -0.1075 | 0.0069 | 392 | -0.1264 | -0.0887 | -15.64 | < 1e-12 | Tukey |

**Supplementary Table 3**

**Table 3a. Quant vs reconstruction**

| Cohort | Platform | Normalization | Contrast | Δz′ | SE | df | 95% CI lower | 95% CI upper | t | Adjusted p | Adjustment |
|---|---|---|---|---|---|---|---|---|---|---|---|
| Mouse | cDNA ONT | CPM | Quantifiers (Kallisto, Oarfish) - reconstruction (Bambu, FLAIR, IsoQuant) | 0.0169 | 0.011 | 1163 | -0.0047 | 0.0385 | 1.53 | 0.126 | None (single test) |
| Mouse | Iso-Seq | TPM | Quantifiers (Kallisto, Oarfish) - reconstruction (Bambu, FLAIR, IsoQuant) | 0.033 | 0.011 | 1163 | 0.0113 | 0.0546 | 2.99 | 0.00285 | None (single test) |
| Mouse | Kinnex | TPM | Quantifiers (Kallisto, Oarfish) - reconstruction (Bambu, FLAIR, IsoQuant) | 0.0431 | 0.0134 | 1163 | 0.017 | 0.0693 | 3.23 | 0.00126 | None (single test) |
| iNeuron | cDNA ONT | CPM | Quantifiers (Kallisto, Oarfish) - reconstruction (Bambu, FLAIR, IsoQuant) | 0.0805 | 0.0066 | 261 | 0.0675 | 0.0935 | 12.16 | < 1e-12 | None (single test) |
| iNeuron | Kinnex | TPM | Quantifiers (Kallisto, Oarfish) - reconstruction (Bambu, FLAIR, IsoQuant) | 0.0365 | 0.0066 | 261 | 0.0235 | 0.0496 | 5.52 | 8.37e-08 | None (single test) |
| LongBench | cDNA ONT | CPM | Quantifiers (Kallisto, Oarfish) - reconstruction (Bambu, FLAIR, IsoQuant) | 0.0926 | 0.0068 | 392 | 0.0792 | 0.1059 | 13.65 | < 1e-12 | None (single test) |
| LongBench | dRNA ONT | CPM | Quantifiers (Kallisto, Oarfish) - reconstruction (Bambu, FLAIR, IsoQuant) | 0.0719 | 0.0068 | 392 | 0.0585 | 0.0852 | 10.6 | < 1e-12 | None (single test) |
| LongBench | Kinnex | TPM | Quantifiers (Kallisto, Oarfish) - reconstruction (Bambu, FLAIR, IsoQuant) | 0.131 | 0.0068 | 392 | 0.1177 | 0.1443 | 19.32 | < 1e-12 | None (single test) |

**Table 3b. Tool pairwise (Tukey)**

| Cohort | Platform | Normalization | Contrast | Δz′ | SE | df | 95% CI lower | 95% CI upper | t | Adjusted p | Adjustment |
|---|---|---|---|---|---|---|---|---|---|---|---|
| Mouse | cDNA ONT | CPM | Bambu - FLAIR | 0.0357 | 0.0171 | 1163 | -0.011 | 0.0824 | 2.09 | 0.227 | Tukey |
| Mouse | cDNA ONT | CPM | Bambu - IsoQuant | 0.0712 | 0.0171 | 1163 | 0.0245 | 0.1179 | 4.17 | 0.000321 | Tukey |
| Mouse | cDNA ONT | CPM | Bambu - Kallisto | 0.0274 | 0.0171 | 1163 | -0.0193 | 0.0741 | 1.6 | 0.495 | Tukey |
| Mouse | cDNA ONT | CPM | Bambu - Oarfish | 0.01 | 0.0171 | 1163 | -0.0367 | 0.0567 | 0.59 | 0.977 | Tukey |
| Mouse | cDNA ONT | CPM | FLAIR - IsoQuant | 0.0355 | 0.0171 | 1163 | -0.0112 | 0.0822 | 2.08 | 0.23 | Tukey |
| Mouse | cDNA ONT | CPM | FLAIR - Kallisto | -0.0083 | 0.0171 | 1163 | -0.055 | 0.0384 | -0.48 | 0.989 | Tukey |
| Mouse | cDNA ONT | CPM | FLAIR - Oarfish | -0.0256 | 0.0171 | 1163 | -0.0723 | 0.0211 | -1.5 | 0.563 | Tukey |
| Mouse | cDNA ONT | CPM | IsoQuant - Kallisto | -0.0438 | 0.0171 | 1163 | -0.0905 | 0.0029 | -2.56 | 0.0781 | Tukey |
| Mouse | cDNA ONT | CPM | IsoQuant - Oarfish | -0.0612 | 0.0171 | 1163 | -0.1079 | -0.0145 | -3.58 | 0.0033 | Tukey |
| Mouse | cDNA ONT | CPM | Kallisto - Oarfish | -0.0174 | 0.0171 | 1163 | -0.0641 | 0.0293 | -1.02 | 0.848 | Tukey |
| Mouse | Iso-Seq | TPM | Bambu - FLAIR | 0.0899 | 0.0171 | 1163 | 0.0432 | 0.1366 | 5.26 | 1.71e-06 | Tukey |
| Mouse | Iso-Seq | TPM | Bambu - IsoQuant | 0.0812 | 0.0171 | 1163 | 0.0345 | 0.1279 | 4.75 | 2.27e-05 | Tukey |
| Mouse | Iso-Seq | TPM | Bambu - Kallisto | 0.056 | 0.0171 | 1163 | 0.0093 | 0.1027 | 3.28 | 0.00952 | Tukey |
| Mouse | Iso-Seq | TPM | Bambu - Oarfish | -0.008 | 0.0171 | 1163 | -0.0547 | 0.0387 | -0.47 | 0.99 | Tukey |
| Mouse | Iso-Seq | TPM | FLAIR - IsoQuant | -0.0087 | 0.0171 | 1163 | -0.0554 | 0.038 | -0.51 | 0.986 | Tukey |
| Mouse | Iso-Seq | TPM | FLAIR - Kallisto | -0.0339 | 0.0171 | 1163 | -0.0806 | 0.0128 | -1.98 | 0.275 | Tukey |
| Mouse | Iso-Seq | TPM | FLAIR - Oarfish | -0.0978 | 0.0171 | 1163 | -0.1445 | -0.0512 | -5.72 | 1.31e-07 | Tukey |
| Mouse | Iso-Seq | TPM | IsoQuant - Kallisto | -0.0251 | 0.0171 | 1163 | -0.0718 | 0.0215 | -1.47 | 0.581 | Tukey |
| Mouse | Iso-Seq | TPM | IsoQuant - Oarfish | -0.0891 | 0.0171 | 1163 | -0.1358 | -0.0424 | -5.21 | 2.17e-06 | Tukey |
| Mouse | Iso-Seq | TPM | Kallisto - Oarfish | -0.064 | 0.0171 | 1163 | -0.1107 | -0.0173 | -3.74 | 0.00178 | Tukey |
| Mouse | Kinnex | TPM | Bambu - FLAIR | 0.0461 | 0.0207 | 1163 | -0.0104 | 0.1026 | 2.23 | 0.17 | Tukey |
| Mouse | Kinnex | TPM | Bambu - IsoQuant | 0.0267 | 0.0207 | 1163 | -0.0298 | 0.0832 | 1.29 | 0.696 | Tukey |
| Mouse | Kinnex | TPM | Bambu - Kallisto | 0.0099 | 0.0207 | 1163 | -0.0467 | 0.0664 | 0.48 | 0.989 | Tukey |
| Mouse | Kinnex | TPM | Bambu - Oarfish | -0.0476 | 0.0207 | 1163 | -0.1041 | 0.0089 | -2.3 | 0.145 | Tukey |
| Mouse | Kinnex | TPM | FLAIR - IsoQuant | -0.0194 | 0.0207 | 1163 | -0.0759 | 0.0372 | -0.94 | 0.883 | Tukey |
| Mouse | Kinnex | TPM | FLAIR - Kallisto | -0.0362 | 0.0207 | 1163 | -0.0927 | 0.0203 | -1.75 | 0.403 | Tukey |
| Mouse | Kinnex | TPM | FLAIR - Oarfish | -0.0937 | 0.0207 | 1163 | -0.1502 | -0.0372 | -4.53 | 6.37e-05 | Tukey |
| Mouse | Kinnex | TPM | IsoQuant - Kallisto | -0.0169 | 0.0207 | 1163 | -0.0734 | 0.0396 | -0.82 | 0.926 | Tukey |
| Mouse | Kinnex | TPM | IsoQuant - Oarfish | -0.0743 | 0.0207 | 1163 | -0.1308 | -0.0178 | -3.59 | 0.00312 | Tukey |
| Mouse | Kinnex | TPM | Kallisto - Oarfish | -0.0575 | 0.0207 | 1163 | -0.114 | -0.001 | -2.78 | 0.0439 | Tukey |
| iNeuron | cDNA ONT | CPM | Bambu - FLAIR | 0.0803 | 0.0103 | 261 | 0.0521 | 0.1084 | 7.82 | 1.38e-12 | Tukey |
| iNeuron | cDNA ONT | CPM | Bambu - IsoQuant | 0.0571 | 0.0103 | 261 | 0.0289 | 0.0852 | 5.56 | 6.54e-07 | Tukey |

| Cohort | Platform | Normalization | Contrast | Δz′ | SE | df | 95% CI lower | 95% CI upper | t | Adjusted p | Adjustment |
|---|---|---|---|---|---|---|---|---|---|---|---|
| iNeuron | cDNA ONT | CPM | Bambu - Kallisto | -0.0621 | 0.0103 | 261 | -0.0903 | -0.034 | -6.06 | 4.73e-08 | Tukey |
| iNeuron | cDNA ONT | CPM | Bambu - Oarfish | -0.0073 | 0.0103 | 261 | -0.0355 | 0.0209 | -0.71 | 0.954 | Tukey |
| iNeuron | cDNA ONT | CPM | FLAIR - IsoQuant | -0.0232 | 0.0103 | 261 | -0.0514 | 0.005 | -2.26 | 0.161 | Tukey |
| iNeuron | cDNA ONT | CPM | FLAIR - Kallisto | -0.1424 | 0.0103 | 261 | -0.1706 | -0.1142 | -13.88 | < 1e-12 | Tukey |
| iNeuron | cDNA ONT | CPM | FLAIR - Oarfish | -0.0875 | 0.0103 | 261 | -0.1157 | -0.0594 | -8.54 | < 1e-12 | Tukey |
| iNeuron | cDNA ONT | CPM | IsoQuant - Kallisto | -0.1192 | 0.0103 | 261 | -0.1474 | -0.091 | -11.62 | < 1e-12 | Tukey |
| iNeuron | cDNA ONT | CPM | IsoQuant - Oarfish | -0.0643 | 0.0103 | 261 | -0.0925 | -0.0362 | -6.27 | 1.45e-08 | Tukey |
| iNeuron | cDNA ONT | CPM | Kallisto - Oarfish | 0.0549 | 0.0103 | 261 | 0.0267 | 0.083 | 5.35 | 1.92e-06 | Tukey |
| iNeuron | Kinnex | TPM | Bambu - FLAIR | 0.084 | 0.0103 | 261 | 0.0558 | 0.1121 | 8.19 | < 1e-12 | Tukey |
| iNeuron | Kinnex | TPM | Bambu - IsoQuant | 0.0534 | 0.0103 | 261 | 0.0252 | 0.0815 | 5.2 | 3.94e-06 | Tukey |
| iNeuron | Kinnex | TPM | Bambu - Kallisto | 0.041 | 0.0103 | 261 | 0.0129 | 0.0692 | 4 | 0.000781 | Tukey |
| iNeuron | Kinnex | TPM | Bambu - Oarfish | -0.0225 | 0.0103 | 261 | -0.0507 | 0.0057 | -2.19 | 0.185 | Tukey |
| iNeuron | Kinnex | TPM | FLAIR - IsoQuant | -0.0306 | 0.0103 | 261 | -0.0588 | -0.0024 | -2.98 | 0.0257 | Tukey |
| iNeuron | Kinnex | TPM | FLAIR - Kallisto | -0.0429 | 0.0103 | 261 | -0.0711 | -0.0148 | -4.19 | 0.000372 | Tukey |
| iNeuron | Kinnex | TPM | FLAIR - Oarfish | -0.1065 | 0.0103 | 261 | -0.1346 | -0.0783 | -10.38 | < 1e-12 | Tukey |
| iNeuron | Kinnex | TPM | IsoQuant - Kallisto | -0.0123 | 0.0103 | 261 | -0.0405 | 0.0158 | -1.2 | 0.75 | Tukey |
| iNeuron | Kinnex | TPM | IsoQuant - Oarfish | -0.0759 | 0.0103 | 261 | -0.104 | -0.0477 | -7.4 | 1.91e-11 | Tukey |
| iNeuron | Kinnex | TPM | Kallisto - Oarfish | -0.0635 | 0.0103 | 261 | -0.0917 | -0.0354 | -6.19 | 2.25e-08 | Tukey |
| LongBench | cDNA ONT | CPM | Bambu - FLAIR | 0.1056 | 0.0105 | 392 | 0.0768 | 0.1344 | 10.05 | < 1e-12 | Tukey |
| LongBench | cDNA ONT | CPM | Bambu - IsoQuant | 0.0577 | 0.0105 | 392 | 0.0289 | 0.0865 | 5.49 | 7.02e-07 | Tukey |
| LongBench | cDNA ONT | CPM | Bambu - Kallisto | -0.0861 | 0.0105 | 392 | -0.1149 | -0.0573 | -8.2 | < 1e-12 | Tukey |
| LongBench | cDNA ONT | CPM | Bambu - Oarfish | 0.0099 | 0.0105 | 392 | -0.0189 | 0.0386 | 0.94 | 0.882 | Tukey |
| LongBench | cDNA ONT | CPM | FLAIR - IsoQuant | -0.0479 | 0.0105 | 392 | -0.0767 | -0.0191 | -4.56 | 6.74e-05 | Tukey |
| LongBench | cDNA ONT | CPM | FLAIR - Kallisto | -0.1917 | 0.0105 | 392 | -0.2205 | -0.1629 | -18.25 | < 1e-12 | Tukey |
| LongBench | cDNA ONT | CPM | FLAIR - Oarfish | -0.0957 | 0.0105 | 392 | -0.1245 | -0.067 | -9.12 | < 1e-12 | Tukey |
| LongBench | cDNA ONT | CPM | IsoQuant - Kallisto | -0.1438 | 0.0105 | 392 | -0.1726 | -0.1151 | -13.69 | < 1e-12 | Tukey |
| LongBench | cDNA ONT | CPM | IsoQuant - Oarfish | -0.0479 | 0.0105 | 392 | -0.0766 | -0.0191 | -4.56 | 6.79e-05 | Tukey |
| LongBench | cDNA ONT | CPM | Kallisto - Oarfish | 0.096 | 0.0105 | 392 | 0.0672 | 0.1248 | 9.14 | < 1e-12 | Tukey |
| LongBench | dRNA ONT | CPM | Bambu - FLAIR | 0.1352 | 0.0105 | 392 | 0.1065 | 0.164 | 12.88 | < 1e-12 | Tukey |
| LongBench | dRNA ONT | CPM | Bambu - IsoQuant | -0.032 | 0.0105 | 392 | -0.0608 | -0.0032 | -3.05 | 0.0207 | Tukey |
| LongBench | dRNA ONT | CPM | Bambu - Kallisto | -0.0709 | 0.0105 | 392 | -0.0997 | -0.0421 | -6.75 | 5.21e-10 | Tukey |
| LongBench | dRNA ONT | CPM | Bambu - Oarfish | -0.004 | 0.0105 | 392 | -0.0328 | 0.0248 | -0.38 | 0.995 | Tukey |
| LongBench | dRNA ONT | CPM | FLAIR - IsoQuant | -0.1672 | 0.0105 | 392 | -0.196 | -0.1385 | -15.92 | < 1e-12 | Tukey |
| LongBench | dRNA ONT | CPM | FLAIR - Kallisto | -0.2061 | 0.0105 | 392 | -0.2349 | -0.1773 | -19.62 | < 1e-12 | Tukey |
| LongBench | dRNA ONT | CPM | FLAIR - Oarfish | -0.1393 | 0.0105 | 392 | -0.168 | -0.1105 | -13.26 | < 1e-12 | Tukey |
| LongBench | dRNA ONT | CPM | IsoQuant - Kallisto | -0.0389 | 0.0105 | 392 | -0.0677 | -0.0101 | -3.7 | 0.00227 | Tukey |
| LongBench | dRNA ONT | CPM | IsoQuant - Oarfish | 0.028 | 0.0105 | 392 | -8.000e-04 | 0.0568 | 2.66 | 0.0613 | Tukey |
| LongBench | dRNA ONT | CPM | Kallisto - Oarfish | 0.0669 | 0.0105 | 392 | 0.0381 | 0.0956 | 6.37 | 5.42e-09 | Tukey |
| LongBench | Kinnex | TPM | Bambu - FLAIR | 0.1278 | 0.0105 | 392 | 0.099 | 0.1566 | 12.17 | < 1e-12 | Tukey |
| LongBench | Kinnex | TPM | Bambu - IsoQuant | 0.2364 | 0.0105 | 392 | 0.2076 | 0.2652 | 22.51 | < 1e-12 | Tukey |
| LongBench | Kinnex | TPM | Bambu - Kallisto | 0.0207 | 0.0105 | 392 | -0.008 | 0.0495 | 1.97 | 0.28 | Tukey |
| LongBench | Kinnex | TPM | Bambu - Oarfish | -0.0399 | 0.0105 | 392 | -0.0687 | -0.0111 | -3.8 | 0.00157 | Tukey |
| LongBench | Kinnex | TPM | FLAIR - IsoQuant | 0.1086 | 0.0105 | 392 | 0.0798 | 0.1374 | 10.34 | < 1e-12 | Tukey |
| LongBench | Kinnex | TPM | FLAIR - Kallisto | -0.1071 | 0.0105 | 392 | -0.1359 | -0.0783 | -10.19 | < 1e-12 | Tukey |

| Cohort | Platform | Normalization | Contrast | Δz′ | SE | df | 95% CI lower | 95% CI upper | t | Adjusted p | Adjustment |
|---|---|---|---|---|---|---|---|---|---|---|---|
| LongBench | Kinnex | TPM | FLAIR - Oarfish | -0.1677 | 0.0105 | 392 | -0.1965 | -0.1389 | -15.97 | < 1e-12 | Tukey |
| LongBench | Kinnex | TPM | IsoQuant - Kallisto | -0.2157 | 0.0105 | 392 | -0.2445 | -0.1869 | -20.53 | < 1e-12 | Tukey |
| LongBench | Kinnex | TPM | IsoQuant - Oarfish | -0.2763 | 0.0105 | 392 | -0.3051 | -0.2475 | -26.31 | < 1e-12 | Tukey |
| LongBench | Kinnex | TPM | Kallisto - Oarfish | -0.0606 | 0.0105 | 392 | -0.0894 | -0.0319 | -5.77 | 1.57e-07 | Tukey |
